# Toward a synthesis of phytoplankton community composition methods for global-scale application

**DOI:** 10.1101/2023.09.07.556589

**Authors:** Sasha J. Kramer, Luis M. Bolaños, Dylan Catlett, Alison P. Chase, Michael J. Behrenfeld, Emmanuel S. Boss, E. Taylor Crockford, Stephen J. Giovannoni, Jason R. Graff, Nils Haëntjens, Lee Karp-Boss, Emily E. Peacock, Collin S. Roesler, Heidi M. Sosik, David A. Siegel

**Affiliations:** Earth Research Institute, University of California Santa Barbara, Santa Barbara CA USA; School of Biosciences, University of Exeter, Exeter UK; Biology Department, Woods Hole Oceanographic Institution, Woods Hole MA USA; Applied Physics Laboratory, University of Washington, Seattle WA USA; Department of Botany & Plant Pathology, Oregon State University, Corvallis OR USA; School of Marine Sciences, University of Maine, Orono ME USA; Department of Microbiology, Oregon State University, Corvallis OR USA; Earth and Oceanographic Science Department, Bowdoin College, Brunswick ME USA

**Keywords:** phytoplankton, pigments, cell imaging, flow cytometry, amplicon sequencing

## Abstract

The composition of the marine phytoplankton community has been shown to impact many biogeochemical processes and marine ecosystem services. A variety of methods exist to characterize phytoplankton community composition (PCC), with varying degrees of taxonomic resolution. Accordingly, the resulting PCC determinations are dependent on the method used. Here, we use surface ocean samples collected in the North Atlantic and North Pacific Oceans to compare high performance liquid chromatography (HPLC) pigment-based PCC to four other methods: quantitative cell imaging, flow cytometry, and 16S and 18S rRNA amplicon sequencing. These methods allow characterization of both prokaryotic and eukaryotic PCC across a wide range of size classes. PCC estimates of many taxa resolved at the class level (e.g., diatoms) show strong positive correlations across methods, while other groups (e.g., dinoflagellates) are not well captured by one or more methods. Since variations in phytoplankton pigment concentrations are related to changes in optical properties, this combined dataset expands the potential scope of ocean color remote sensing by associating PCC at the genus- and species-level with group- or class-level PCC from pigments. Quantifying the strengths and limitations of pigment-based PCC methods compared to PCC assessments from amplicon sequencing, imaging, and cytometry methods is the first step toward the robust validation of remote sensing approaches to quantify PCC from space.

## 1. Introduction

Phytoplankton encompass tens of thousands of species and their composition varies broadly across spatiotemporal scales (e.g., Caron et al., 2012; de Vargas et al., 2015). The vast diversity of phytoplankton structures marine food webs, impacts biogeochemical cycling of nutrients, and influences the magnitude of carbon sequestration in the deep ocean by the biological pump (Martiny et al., 2013; Guidi et al., 2016). Phytoplankton diversity also impacts the flux of particulate organic carbon to depth and its vertical remineralization length scale, both of which are important controls on the efficiency of the biological pump (Guidi et al., 2015; Trudnowska et al., 2021; Durkin et al., 2022; Siegel et al., 2023). Furthermore, phytoplankton diversity is correlated with ecosystem productivity and resilience (e.g., Behrenfeld, 2014; Vallina et al., 2017). Quantifying surface ocean phytoplankton community composition (PCC) is essential for a complete understanding of present-day marine ecosystems and the biological pump, and for forecasting future changes in the ecosystem services provided by phytoplankton.

Many methods exist to characterize PCC from field samples, with varying taxonomic resolution, quality control and standardization criteria, and scales of observation (Sosik et al., 2014; Johnson and Martiny, 2015; Lombard et al., 2019). Common methods include microscopy (Karlson et al., 2010), high performance liquid chromatography (HPLC) pigments (e.g., Mackey et al., 1996; Uitz et al., 2006; Kramer and Siegel, 2019), flow cytometry (FCM; e.g., Zubkov et al., 1998; Sosik et al., 2010), quantitative cell imaging (e.g., with the Imaging FlowCytobot [IFCB]; Olson and Sosik, 2007), and amplicon sequencing of “barcode” genes (e.g., Needham and Fuhrman, 2016; Catlett et al., 2020). This list is not exhaustive and does not include optical proxy methods developed for use with in situ and remote sensing approaches (Thibodeau et al., 2014; Uitz et al., 2015). Typically, the appropriate method for targeting PCC relates to the goals of a given study. For instance, approaches that require broad spatial coverage often rely on ocean color methods from satellite remote sensing data to cover the necessary scales (Bracher et al., 2017 and references therein). Alternately, approaches that require high taxonomic resolution favor methods that provide genus- to species-level characterization of PCC, such as amplicon sequencing (Sommeria-Klein et al., 2021).

HPLC pigments are widely used for creating and validating ocean color remote sensing algorithms. HPLC measurements are widespread in the global surface ocean (Kramer and Siegel, 2019), quality-controlled (Hooker et al., 2012), and have clear links to satellite ocean color observations due to the direct impact of phytoplankton pigments on the spectral shape and magnitude of light absorption, and thus remote sensing reflectance (Chase et al., 2013, 2017; Kramer et al., 2022). However, HPLC pigments have significant limitations in describing PCC. The maximum number of groups identified by HPLC pigments depends on the dataset and scale of observation, with typically between 4 and 7 distinct groups separated by a given HPLC dataset (Catlett and Siegel, 2018; Kramer and Siegel, 2019; Kramer et al., 2020a). There are also several caveats to pigment-based taxonomy. Pigment concentration and composition can be affected by light history and nutrient limitation (Schlüter et al., 2000; Henriksen et al., 2002). Species or even strains within a single species can have varying pigment compositions (Zapata et al., 2004; Neeley et al., 2022). Most notably, nearly all phytoplankton groups share some accessory pigments due to their evolutionary history or their feeding strategies (or both), leading to similarities in pigment composition that make statistical chemotaxonomic methods that assume independence between pigments, such as the widely-applied CHEMTAX approach, invalid for assessing PCC (Jeffrey et al., 2011; Catlett and Siegel, 2018; Kramer et al., 2019).

Given the widespread use of HPLC pigments for ocean color PCC algorithm development and validation, it is important to characterize and quantify the information content of HPLC pigments compared to other methods without these same limitations. Here, a dataset of surface ocean HPLC pigment samples is compared to PCC assessments from 18S and 16S rRNA gene amplicon sequencing, quantitative imaging from IFCB, and FCM. Each PCC method has strengths and weaknesses (Table 1; Johnson and Martiny, 2015; Lombard et al., 2019). For instance, nearly all methods capture only part of the phytoplankton community size range, as determined by filter pore size, volume of seawater sampled, and/or resolution of the instrument. Similarly, each method has a (quantifiable or unquantifiable) fraction of “unknown” or “unidentified” phytoplankton. For instance, 16S SSU rRNA gene amplicon sequencing (hereafter, 16S) cannot reliably identify dinoflagellate plastids (Lin et al., 2011) and prokaryotic plankton do not have the 18S SSU rRNA gene (hereafter, 18S). Most smaller cells (<5µm) are unmeasured or unidentifiable by the IFCB due to image resolution and detection sensitivity (Sosik and Olson, 2007), while FCM is limited to separating broad groups of cells (∼1-65 µm). Some of these properties are not inherent to the measurement but change as new iterations of the methods are introduced. For instance, the coverage of phytoplankton diversity by rRNA gene sequencing (both 16S and 18S) is subject to biases caused by natural variations in DNA sequences at the primer-binding sites, which have been continually refined and improved upon by expanded sequencing and by new primer designs.

**Table 1.** Summary of the five PCC methods presented here. For each method, a short overview is provided of the targeted taxonomic range and resolution, the approximate size range captured by the method (*all assume nominal pore sizes; HPLC pigment size range for combusted GF/F filters), the exact measurement provided by each method, and known method strengths and weaknesses. For all methods, the upper size range is influenced by the volume sampled.

| Method | Kingdom | Size range | Taxonomic resolution | Measurement result | Notable strengths | Assorted limitations and challenges |
| --- | --- | --- | --- | --- | --- | --- |
| HPLC pigments | Prokaryotes and eukaryotes | >0.3 $\mu\text{m}^*$ | Group level (dataset dependent) | Pigment concentrations | Direct links to optical properties; publicly available data with global coverage; highly standardized method | Inter- and intra-group pigment variation; environmental impacts on concentration; limited taxonomic resolution |
| 18S amplicon sequencing | Eukaryotes | >0.22 $\mu\text{m}^*$ | Class to species level (dependent on target gene, amplicon length, taxonomic assignment) | Amplicon sequence variant counts | Consistent, high-resolution results for many eukaryotic taxa; publicly available data with global coverage; some approaches for standardizing in development | Gene copy numbers are not equal across taxa; method differences can bias results; attribution of ASVs to taxa is incomplete; no photosynthetic prokaryotes; ASV counts are compositional, not absolute |
| 16S amplicon sequencing | Prokaryotes and many eukaryotes | >0.22 $\mu\text{m}^*$ | Class to species level (dependent on target gene, amplicon length, taxonomic assignment) | Amplicon sequence variant counts | Targets both prokaryotes and eukaryotes at relatively high resolution; publicly available data with global coverage; some approaches for standardizing in development | Cannot identify dinoflagellates; method differences can bias results; attribution of ASVs to taxa is incomplete; gene copy numbers are not equal across taxa; ASV counts are compositional, not absolute |
| Quantitative imaging (IFCB) | Eukaryotes, some colonial prokaryotes | $\sim 6\text{-}150 \mu\text{m}$ | Class to species level (dependent on taxonomic assignment) | Cell counts and biovolumes | Biovolume relates more easily to carbon; cell-level taxonomy (allows for gene- or pigment-per-biovolume); iterative taxonomic ID possible | High fraction of unidentified cells, particularly at low end of size range; can not quantify smaller cells; small volume sampled can select against rare, large cells |
| Flow cytometry | Prokaryotes and eukaryotes | $\sim 0.5\text{-}64 \mu\text{m}$ | Four groups only: *Prochlorococcus*, *Synechococcus*, nano and picoeukaryotes | Cell counts | Cell-level measurements for prokaryotes and some eukaryotes; carbon estimates possible with some assumptions; highly standardized method for groups targeted | Large fractions of unidentified cells (eukaryotes); limited size range; very small volume sampled; quantify cells but cannot accurately capture shape/biovolume |

The strength of each method to describe the “abundance” or “biomass” of a given phytoplankton group can be assessed by both absolute (i.e., cell counts, cell biovolume concentrations) and relative (i.e., relative pigment concentrations, relative amplicon sequence abundances) metrics. The assumptions inherent in some methods limit the interpretation of the results, such as the challenge of unequal copy numbers of the 16S and 18S genes across taxa (e.g., Godhe et al., 2008; de Vargas et al., 2015; Needham and Fuhrman, 2016). Comparisons between and among PCC methods are relatively rare, and reveal variability when different methods are compared (e.g., Not et al., 2008; Couple et al., 2015; Gong et al., 2020; Campbell et al., 2022; Chase et al., 2022; Catlett et al., 2022; Nardelli et al., 2023). In one example, amplicon sequencing and light microscopy each provide high resolution taxonomic information for larger phytoplankton, but abundance patterns did not agree in genus- to species-level comparisons (Abad et al., 2016). In another example, phytoplankton pigment concentrations correlate with relative abundances of amplicon sequencing data for some groups (e.g., cryptophytes) but not for others (e.g., diatoms; Lin et al., 2019). While method comparison often highlight aspects of agreement, differences between methods can also be useful to highlight limitations, strengths, and weaknesses, and can reveal novel insights into microbial ecology (Catlett et al., 2022).

Here, we compared PCC among methods on samples collected in the western North Atlantic as part of the North Atlantic Aerosols and Marine Ecosystems Study (NAAMES; Behrenfeld et al., 2019, 2021) and in the Subarctic North Pacific as part of the EXport Processes in the Ocean from RemoTe Sensing (EXPORTS; Siegel et al., 2021) field campaigns.

Combining two oceanographic regions and multiple PCC methods with diverse measurement strengths and limitations allows for an evaluation of pigment-based PCC assessments relative to other, higher-resolution methods. Our analysis highlights the importance of integrating PCC methods to extend phytoplankton community information beyond the capabilities provided by one method alone.

## 2. Materials and Procedures

Near-surface samples were selected for this analysis to maximize the number of comparisons among the five PCC methods considered. However, the analysis was performed separately across two datasets, each with three methods available to assess PCC, to maximize the number of observations available for comparison. The first dataset focuses on PCC metrics for eukaryotic phytoplankton: HPLC pigments, 18S amplicon sequencing, and quantitative cell imaging by IFCB. This dataset includes 45 samples in total. Twenty-four of these samples were collected in the eastern North Pacific Ocean in August-September 2018 as part of EXPORTS (Figure S1A), where each sampling site has collocated HPLC, 18S, and IFCB data. The remaining 21 samples were collected in the western North Atlantic Ocean in May-June 2016, August-September 2017, and March-April 2018 as part of NAAMES (Figure S1B), where each sampling site has collocated HPLC and 18S data, and 18 sites also include IFCB data.

The second dataset compares PCC metrics for prokaryotic and eukaryotic phytoplankton from HPLC pigments, 16S amplicon sequencing, and cell counts from FCM. This dataset includes 65 concurrent HPLC and 16S samples, 34 of which have coincident FCM samples. All samples were collected in the western North Atlantic Ocean as part of NAAMES, in November 2015, May-June 2016, August-September 2017, and March-April 2018 (Figure S1C).

It is important to note that both HPLC pigment concentrations and identified cell abundances can be compared to other metrics in absolute terms or as relative compositions if normalized to the total pigment concentration or number of cells in the sample. However, for the 16S and 18S amplicon sequencing methods employed here, only relative data are available, as the total number of sequence counts for a given sample or sequencing run are influenced by the sample analysis procedures (Lin, 2011; Gloor et al., 2017; Caron and Hu, 2019). We therefore refer to 16S and 18S amplicon results as “relative sequence abundances” throughout.

### 2.1 HPLC phytoplankton pigments

Surface water samples for HPLC pigment analysis were collected either from Niskin bottles on a CTD rosette or from the ship’s underway flow-through system (≤5 m depth), which used a diaphragm pump to minimize impacts on cells (Cetinić et al., 2016). Two-liter whole seawater samples were filtered onto pre-combusted (450°C for 4 hours) 25 mm Whatman ® GF/F filters. After combustion, the filter pore size has been estimated to be ∼0.3 µm (Nayar and Chou, 2003). Filters were stored in foil packets and frozen in liquid nitrogen immediately after sampling and then kept in liquid nitrogen or at −80°C until analysis. HPLC samples were processed at the NASA Goddard Space Flight Center following Van Heukelem and Hooker (2011) and Hooker et al. (2012).

Degradation pigments (chlorophyllide, phaeophytin, and phaeophorbide) and accessory pigments with limited distinct taxonomic utility (monovinyl chlorophyll-*a*, total chlorophyll b, total chlorophyll c, alpha-beta carotene, diatoxanthin, and diadinoxanthin) were removed from our analysis following Kramer and Siegel (2019). Lutein (an accessory pigment in green algae) was also not considered since it was below detection in >80% of the samples in this dataset.

Concentrations of the remaining 15 pigments were used in this analysis. These pigments are total chlorophyll-*a* (Tchla, a sum of monovinyl chlorophyll-*a*, divinyl chlorophyll a, chlorophyllide, and assorted chlorophyll-*a* allomers and epimers), 19’-hexanoyloxyfucoxanthin (19HexFuco), 19’-butanoyloxyfucoxanthin (19ButFuco), alloxanthin (Allo), fucoxanthin (Fuco), peridinin (Perid), zeaxanthin (Zea), divinyl chlorophyll a (DVchla), monovinyl chlorophyll b (MVchlb), divinyl chlorophyll b (DVchlb), chlorophyll c_1_+c_2_ (Chlc12), chlorophyll c_3_ (Chlc3), neoxanthin (Neo), violaxanthin (Viola), and prasinoxanthin (Pras). Pigment values below the NASA Ocean Biology Processing Group method detection limits (Van Heukelem and Thomas, 2001) were set to zero.

Most accessory pigments are shared between phytoplankton groups (Jeffrey et al., 2011 and references therein), making “biomarker” pigments imprecise identifiers for taxonomy. However, some pigments are used as biomarkers throughout the literature despite extensive documentation that these pigments are not limited to one taxonomic group (e.g., Fuco for diatoms; e.g., Jeffrey et al., 2011 and references therein; Chase et al., 2020, 2022). In the presentation to follow, we used commonly-applied pigment-based taxonomic designations to compare these biomarker approaches to other, higher-resolution methods. These designations are as follows: Fuco (diatoms), Perid (dinoflagellates), 19HexFuco (prymnesiophytes), 19ButFuco (dictyochophytes, pelagophytes), Allo (cryptophytes), DVchla (*Prochlorococcus*), Zea (other cyanobacteria), MVchlb (chlorophytes). The ratios of these accessory pigments to Tchla were used to create phytoplankton composition metrics for comparison with other PCC methods (Table 2), with a goal of assessing the degree of correspondence between pigment-based PCC and higher taxonomic resolution observations. It should be noted here that conventional HPLC methods do not measure some pigments that can be used for taxonomic classification (e.g., phycobilin pigments found in cyanobacteria).

**Table 2.** Major phytoplankton groups addressed in this analysis and their corresponding metrics from biomarker pigments, amplicon sequencing methods, and flow cytometry methods. Note that in many cases, biomarker pigments are not exclusively found in the listed phytoplankton classes and groups (see tables in Catlett and Siegel, 2018 and Kramer and Siegel, 2019 for more detail). The colors used to designate the phytoplankton group in column 1 are consistent with the colors used in other figures and tables throughout this work. *Chlorarachniophyceae are Rhizaria that contain MVchlb. For the purposes of this analysis, they are grouped with other MVchlb-containing taxa.

| Pigment-based phytoplankton group | Pigment | 18S class(es) | 16S class(es) | IFCB group | FCM group |
| --- | --- | --- | --- | --- | --- |
| Diatoms | Fuco | Bacillariophyceae | Bacillariophyceae | Diatoms | N/A |
| *Prochlorococcus* sp. | DVchl*a* | N/A | *Prochlorococcus* | N/A | *Prochlorococcus* |
| Cyanobacteria | Zea | N/A | *Synechococcus* | N/A in this analysis | *Synechococcus* |
| Dinoflagellates | Perid | Dinophyceae | N/A | Dinoflagellates | N/A |
| Dictyochophytes, Pelagophytes | 19ButFuco | Dictyochophyceae, Pelagophyceae | Dictyochophyceae, Pelagophyceae | Dictyochophytes | N/A |
| Prymnesiophytes (haptophytes) | 19HexFuco | Prymnesiophyceae | Prymnesiophyceae, Rappemonad | Prymnesiophytes | N/A |
| Green algae | MVchl*b* | Chlorarachniophyceae*, Chloropicophyceae, Mamiellophyceae, Pyramimonadophyceae | Bathycoccus, Micromonas, Ostreococcus, Prasinophyceae | Chlorophytes | N/A |
| Cryptophytes | Allo | Cryptophyceae | Cryptophyceae | Cryptophytes | N/A |

### 2.2 16S amplicon sequencing

Samples for 16S amplicon sequencing were collected at the same time as HPLC pigment samples on NAAMES, either from the flow-through system or from Niskin bottle sampling. Detailed methodology for sample collection and preparation can be found in Bolaños et al. (2020) and (2021). In brief, each whole seawater sample was filtered onto a Sterivex filter with a 0.22 µm pore size, 1 mL of sucrose lysis buffer was added to the filter, and then filters were stored at −80°C until further processing. The methods used here targeted the V1-V2 region of the 16S rRNA gene. All samples were prepared following a standard Illumina 16S sequencing preparation protocol. Sequencing was conducted at the Center for Quantitative Life Sciences (Oregon State University, Corvallis, OR USA).

Sequences were trimmed, amplicon sequence variants (ASVs) were determined, and chimeras were removed with the DADA2 (v. 1.2) package for R (Callahan et al., 2016). Taxonomy was then assigned to sequences with the assignTaxonomy command in DADA2 and the SILVA gene database (v. 123; Quast et al., 2012; Yilmaz et al., 2014). Taxonomy was also assigned and confirmed from phylogenetic tree placement via Phyloassigner (v. 089; Vergin et al., 2013). The 1594 resulting phytoplankton and bacterial ASVs were then condensed into 45 phytoplankton groups. Fourteen of those groups were >1% abundant in at least one of the 65 matchup samples and subsequently were used in analyses. These taxonomic groups included *Prochlorococcus* sp., *Synechococcus* sp., Bacillariophyceae (diatoms), Bolidophyceae, Chrysophyceae, Prymnesiophyceae, Rappemonads, Dictyochophyceae, Pelagophyceae, Cryptophyceae, *Bathycoccus* sp. (chlorophyte), *Micromonas* sp. (chlorophyte), *Ostreococcus* sp. (chlorophyte), and Prasinophyceae (chlorophyte). 16S amplicon sequencing detects many prokaryotic and chloroplast-containing eukaryotic taxa, but notably does not capture photosynthetic dinoflagellates, which have acquired plastids relatively recently in evolutionary history via successive endosymbioses (Lin, 2011; Table 2). All 16S data considered here were examined in compositional space. 16S copy numbers tend to vary less than some other genes (Needham and Fuhrman, 2016) and instead tend to covary with the number of chloroplast genomes per cell, which can impact comparisons to other methods.

### 2.3 18S amplicon sequencing

All 18S amplicon sequencing samples from NAAMES and EXPORTS were collected concurrently with surface HPLC samples. The NAAMES 18S amplicon samples (N = 21) were sequenced from DNA extracted for the 16S samples. The EXPORTS 18S samples (N = 24) were collected similarly to the NAAMES samples. Specifically, whole seawater samples were collected from the flow-through system and filtered on Sterivex filters with a 0.22 µm pore size at low pressure. One mL sucrose lysis buffer was added to each filter before storing at −80°C. Subsequent processing targeted the V9 region of the 18S gene. All samples were prepared following the methods presented in Catlett et al. (2020). Samples were sequenced in three batches between July 2020 and December 2020. Each batch included negative control and mock community positive control samples (Catlett et al., 2020) to ensure consistency between sequencing runs. Sequencing was conducted with a MiSeq PE150 v2 kit (Illumina) at the DNA Technologies Core of the University of California Davis Genome Center (Davis, CA USA).

The DADA2 (v. 1.12) package was used to trim sequences, infer ASVs, and remove chimeras. Taxonomy was assigned to ASVs with the ensembleTax method developed by Catlett et al. (2021a), which combines the results of the assignTaxonomy function in the DADA2 pipeline (Callahan et al., 2016) with the results of the IDTAXA function from the DECIPHER Bioconductor package (v. 2.2; Murali et al., 2018) and considers both the Protist Ribosomal Reference (PR2; v. 4.14; Guillou et al., 2012) and SILVA (v. 138; Quast et al., 2012; Yilmaz et al., 2014) databases. The PR2 taxonomic nomenclature is used below. ensembleTax results in a collection of relatively high-resolution taxonomic assignments for each ASV. All ASVs of non-protistan origin were removed (Catlett et al., 2022), leaving 2433 unique ASVs. Phytoplanktonic ASVs were then separated from other protists after assigning putative feeding strategies based on the ensemble taxonomy predictions for each ASV (Catlett et al., 2022).

Of the 2433 protistan ASVs, 635 were identified as phytoplankton. ASVs were then aggregated to the class level to consider classes with >1% abundance in any sample. The present analysis focuses on those 13 classes, comprised of 135 ASVs: Bacillariophyceae (diatoms), Dinophyceae (dinoflagellates), Bolidophyceae, Chrysophyceae, MOCH-2 (red algae), Prymnesiophyceae, Dictyochophyceae, Pelagophyceae, Cryptophyceae, Chloroarachniophyceae (MVchlb-containing Rhizaria), Chloropicophyceae (chlorophyte), Mamiellophyceae (chlorophyte), and Pyramimonadophyceae (chlorophyte). While 18S reliably separates many eukaryotes, this gene is not found in prokaryotes (Table 2).

Because total sequence counts vary based on methodology and sample processing approaches, all 18S data considered here are compared to other methods in relative space. 18S relative sequence abundances scale with cell size for many taxa and often show good qualitative agreement with biomass fractions from other methods (Zhu et al., 2005; Godhe et al., 2008; de Vargas et al., 2015).

### 2.4 Quantitative cell imaging (IFCB)

During both the NAAMES and EXPORTS field campaigns, an IFCB (McLane Research Laboratories, Inc.) was used to evaluate community composition in samples from the ship’s flow-through system (intake ≤5m). IFCB was configured to analyze a new 5-ml sample taken automatically every 20-25 minutes. Precise sample volume varies as a function of cell concentration and the volume is recorded to allow for calculation of quantitative cell concentrations (Olson and Sosik, 2007). Matched samples were selected based on the time and location of sample collection (<0.1 degree latitude or longitude apart, ±2 hours apart). If multiple IFCB samples were collected within the hour of and at the same location (based on latitude and longitude) as discrete sample collection (HPLC pigments, 18S amplicon sequencing), then multiple (up to 3) IFCB samples were aggregated to create one matchup sample. IFCB imaged all cells and particles (∼6-150 µm diameter) that triggered a signal above a defined threshold in fluorescence or scattering (Olson and Sosik, 2007; Haëntjens et al., 2022). These images were automatically classified, followed by manual verification and error correction as described below. For both field campaigns, cell biovolume concentrations were estimated following Moberg and Sosik (2012) and updates to that method (https://github.com/hsosik/ifcb-analysis/tree/features_v3).

Detailed methodology for the taxonomic assignment of IFCB imagery on NAAMES can be found in Chase et al. (2020). In summary, the 250,660 images used here were exported to the web platform EcoTaxa (Picheral et al., 2017). A supervised random forest machine learning approach was used to predict the classification of each image into 84 pre-determined sets, and the automated classification was confirmed or corrected manually. Non-living and detrital particles were separated from living cells, and living cells were annotated with the most detailed taxonomic designation possible. Following the automated and manual classification and validation, the diversity of living phytoplankton cells was condensed into seven taxonomic categories selected to match the pigment-based phytoplankton groups as closely as possible: diatoms, dinoflagellates, dictyochophytes, prymnesiophytes, cryptophytes, euglenoids, chlorophytes, and “other” (which includes unidentifiable living cells, some of which may potentially belong to one of those seven categories, as well as all other taxonomic groups not described by the prior categories).

The EXPORTS images were automatically classified with a supervised convolutional neural network of Inception v3 architecture (Szegedy et al., 2015). This network was initialized with pre-trained weights from ImageNet (Russakovsky et al., 2015) and then fine-tuned with a 49-category training set of IFCB images. Application of the trained model allowed us to separate This machine learning approach separated 177,161 images into the 49 pre-determined categories, including detritus or non-phytoplankton (which were removed from further analysis) and many classes of living phytoplankton cells. The results of the automated classifier were confirmed or corrected via sequential (2x) manual verification. Once all images were classified and validated, the EXPORTS images were aggregated into the same seven groups as the NAAMES dataset. There were no euglenoids or green algae identified in the EXPORTS matchup dataset. However, these groups are still included for comparison. As configured on these cruises, IFCBs did not comprehensively image cells smaller than ∼6 µm diameter; hence, most small nano- and all pico-phytoplankton are not assessed.

### 2.5 Flow cytometry (FCM)

Full methodological details of flow cytometric analysis on NAAMES can be found in Graff and Behrenfeld (2018). Briefly, flow cytometry was performed by a calibrated BD Influx Cell Sorter (ICS) on whole, unpreserved surface seawater samples collected from Niskin bottles and from the ship’s flow-through system (≤5 m). In each sample, a minimum of 7,000 total cells were interrogated. The counts per sample were transformed into cell concentrations based on calculated sample flow rates (Graff et al., 2018). Data were classified into a mixture of four taxonomic and size-based categories: *Prochlorococcus* sp., *Synechococcus* sp., picoeukaryotes, and nanoeukaryotes (limited to diameters ≤64 µm, determined in lab and at sea from cultures). While some micro-sized eukaryotes are included in the definition of nanoeukaryotes from FCM, those cells were not large contributors to this dataset (Haëntjens et al., 2022). The FCM-derived groups were defined by the scattering and fluorescence properties associated with each category, which allows cells to be separated from one another. As with the IFCB samples, matchups between flow cytometry and other discrete samples were defined by collocation in space and time. A matchup sample was defined if FCM samples were collected in the same place (e.g., same latitude and longitude) within ±2 hours of concurrent HPLC and 16S amplicon sequencing samples. Nano- and micro-sized phytoplankton are not taxonomically separable by FCM.

### 2.6 Environmental data

Environmental data recorded during NAAMES and EXPORTS included determinations of sea surface temperature (SST), salinity, mixed layer depth (MLD), and incident photosynthetically active radiation (PAR). All environmental data were matched with the closest PCC sample in space and time (with no matches greater than 15 minutes apart). SST and salinity were collected from the ships’ underway systems. MLD was calculated for all samples where there were coincident CTD profiles (details in Della Penna and Gaube [2019] for NAAMES and Siegel et al. [2021] for EXPORTS). Finally, PAR was measured with a LICOR cosine sensor, mounted to avoid the impact of ship shadow as much as possible (further details available on NASA’s SeaBASS repository for both field campaigns: https://seabass.gsfc.nasa.gov/archive/OSU/NAAMES/ and https://seabass.gsfc.nasa.gov/archive/OSU/behrenfeld/EXPORTS/EXPORTSNP). The average surface PAR value for the 24 hours prior to each HPLC sample was used to represent the time scale relevant to cell physiology and pigment production.

### 2.7 Statistical methods

Correlation matrices were constructed following Kramer et al. (2020a), where correlations between variables (relative abundances of phytoplankton taxa) were weighted following the Weighted Gene Co-Expression Network Analysis (Zhang and Horvath, 2005) to maximize within-group correlations and minimize between-group correlations, thereby highlighting the strongest connections between methods. Chord diagrams (Gu et al., 2014) were constructed with the “circlize” package in R (v. 4.1.2) applied to the weighted correlation matrices among pigments and PCC metrics for relative sequence abundances aggregated to the most abundant group for both (1) 18S and relative cell biovolume concentrations from the IFCB and (2) 16S and relative cell counts from FCM. A network graph was constructed from the weighted correlation matrices with the “graph” function in MATLAB. Variables were colored by the results of a network-based community detection analysis following Kramer et al. (2020a), with the “modularity_und” function for MATLAB (Rubinov and Sporns, 2010; Brain Connectivity Toolbox, https://sites.google.com/site/bctnet/Home). Notably, all analyses are limited by the uncertainties associated with each PCC method, which can be large for some approaches (see Chase et al., 2022).

## 3. Assessment

### 3.1 Trends in PCC from HPLC pigments, 18S amplicons, and IFCB cell biovolume concentrations

Our comparison of pigment-based PCC to other methods across the aggregated eukaryotic dataset reveals varied relationships across taxonomic groups (Figures 1 and 2; Table 2). Median Fuco concentrations, diatom relative sequence abundance, and diatom biovolume concentrations are consistently high across all three methods and across cruises (Figures 1 and 2). While median Perid concentrations are low compared to other accessory pigments (and lower on EXPORTS than NAAMES; Figure 1A-B), median dinoflagellate relative sequence abundances and dinoflagellate biovolume concentrations are high for all samples (Figure 1C-F). We observe that median 19HexFuco concentrations are relatively high compared to other accessory pigment concentrations, particularly during the NAAMES campaigns (Figure 1A), which is consistent with high prymnesiophyte relative sequence abundances (Figure 1C) but not with biovolume estimates from the IFCB, which show relatively lower median prymnesiophyte biovolume concentrations compared to other groups measured by the IFCB (Figure 1E).

**Figure 1.**
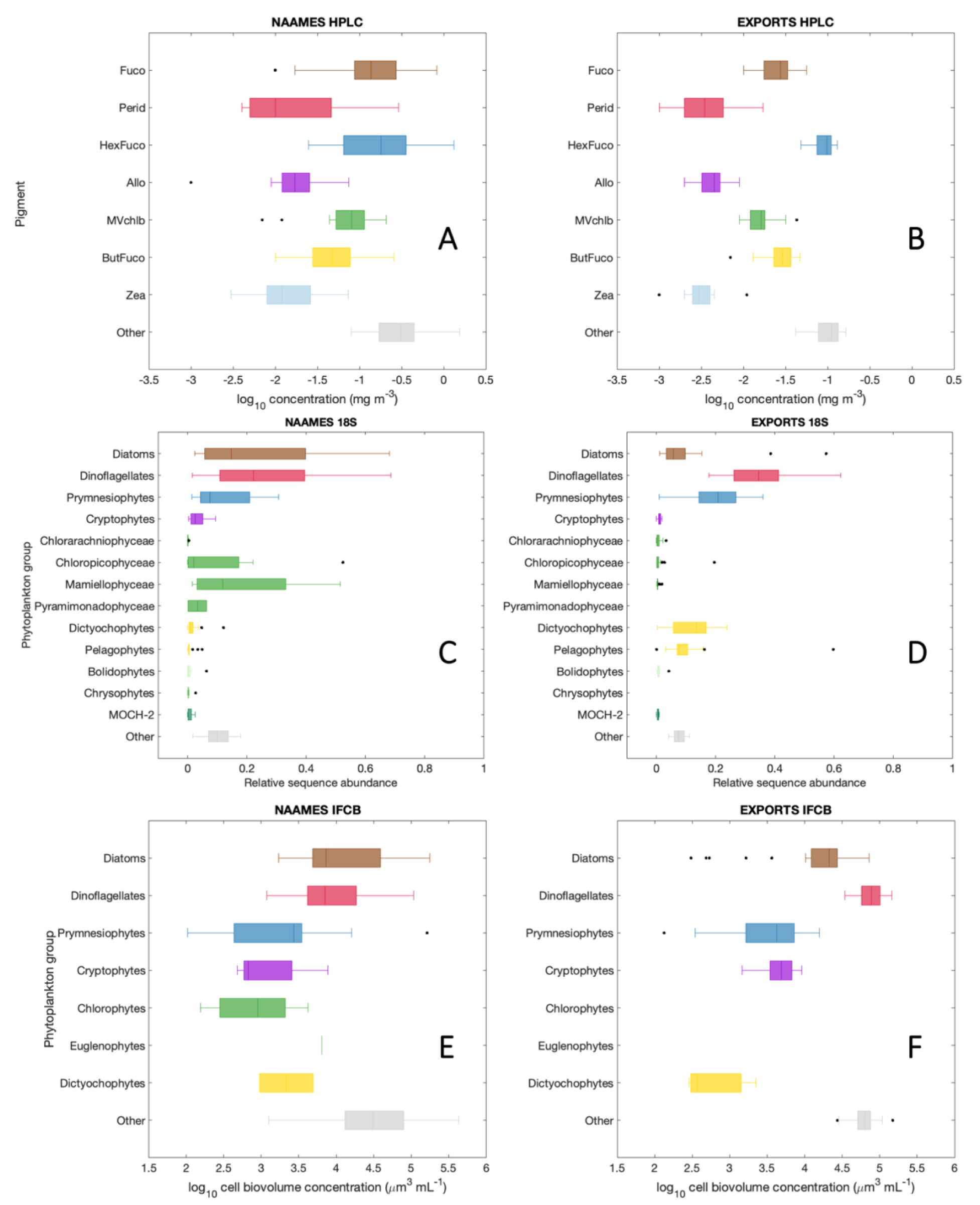
Pigment concentrations measured on (A) NAAMES and (B) EXPORTS; 18S relative sequence abundances on (C) NAAMES and (D) EXPORTS; and cell biovolume concentrations on (E) NAAMES and (F) EXPORTS. The box shows the median value and encompasses the upper and lower quartiles; whiskers span the non-outlier minimum and maximum values; outliers (black dots) are any samples that fall greater than 1.5x the interquartile range from the top or bottom of the box. Boxes are colored similarly for shared groups: diatoms in brown, prymnesiophytes in dark blue, cryptophytes in purple, chlorophytes in bright green, and dictyochophytes + pelagophytes in gold. Grey boxes indicate the “other” fraction for each group.

**Figure 2.**
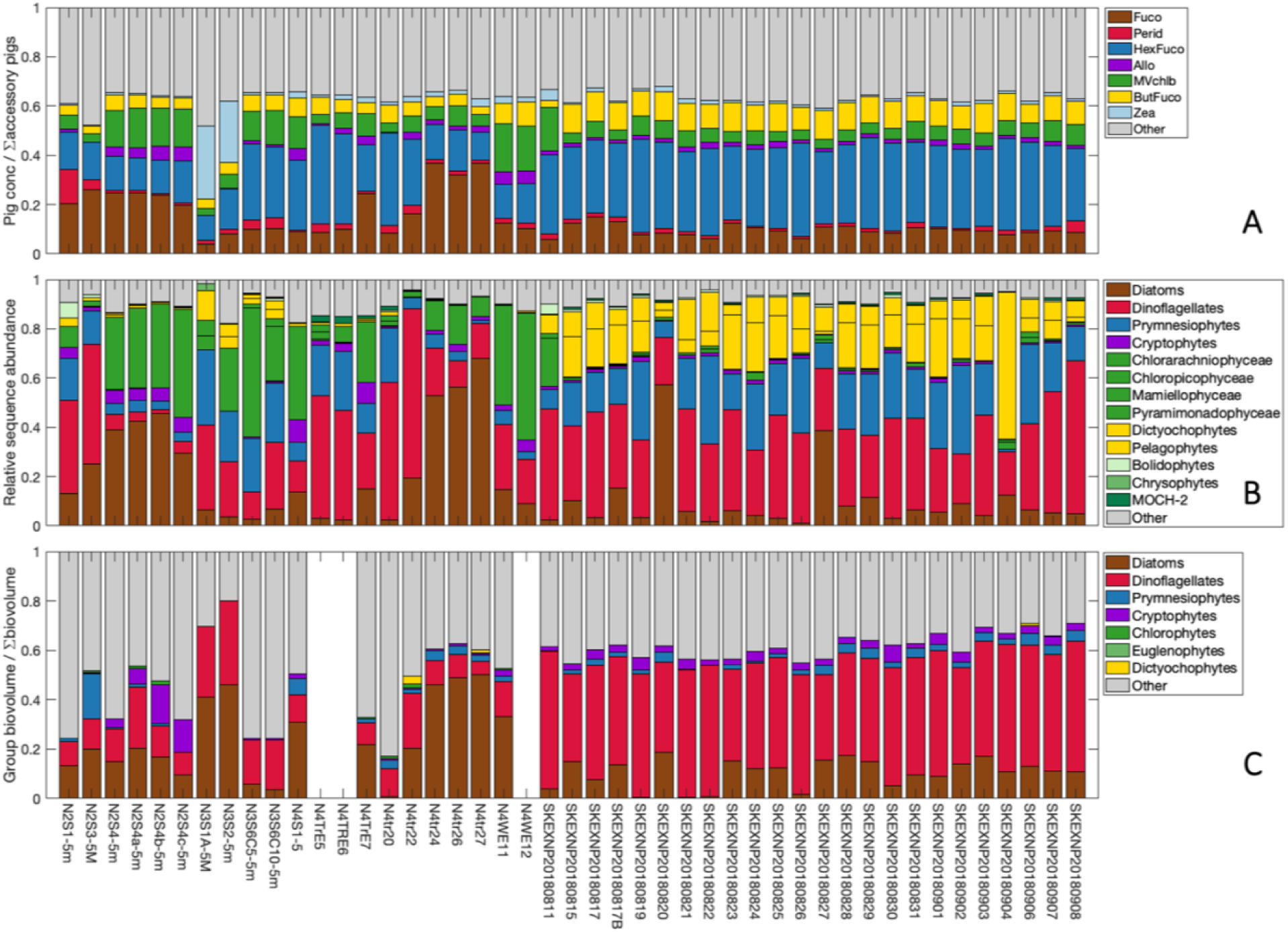
Relative fractions of (A) phytoplankton pigments to summed accessory pigments; (B) 18S sequences; and (C) IFCB biovolume concentrations from NAAMES and EXPORTS. Samples are organized from left to right in the order collected, from NAAMES 2-4 on the left half and EXPORTS on the right half. Bars are colored similarly for shared groups: diatoms in brown, prymnesiophytes in dark blue, cryptophytes in purple, chlorophytes in bright green, and dictyochophytes + pelagophytes in gold. Grey bars indicate the “other” fraction for each group.

Similarly, there are consistent fractions of cryptophyte markers across datasets (2-4%), including median relative Allo concentrations, relative cryptophyte sequences, and relative cryptophyte biovolume concentrations. When cryptophytes are absent, they are absent across all methods. Median 19ButFuco concentrations are similar between NAAMES and EXPORTS (Figure 1A-B), but dictyochophyte and pelagophyte relative sequence abundances are much higher during EXPORTS than NAAMES (Figure 1C-D). There were very few dictyochophytes observed in the EXPORTS IFCB imagery, with higher dictyochophyte biovolume concentrations in the NAAMES data (Figure 1E-F). The relative fraction of “other” accessory pigments and “other” cell biovolume concentrations is higher than the relative fraction of “other” sequences.

Rather than as a composite for the dataset as a whole, the compositional trends described above can also be viewed across samples (Figure 2). For each of the three PCC methods, the phytoplankton community is much more consistent between samples during EXPORTS than NAAMES, which is expected given the broader spatiotemporal range of the NAAMES sampling. Pigment concentrations were normalized to the sum of all accessory pigments (which is highly correlated with Tchla in this dataset; R^2^ = 0.96 and hereafter referred to as ∑ *pigs*) for comparison with the other two methods. Perid ratios to summed accessory pigments are notably lower than relative dinoflagellate sequence abundance, which are in turn lower than relative dinoflagellate biovolume concentrations determined by IFCB. In contrast, the 19HexFuco/∑ *pigs* ratio is always greater than the relative fraction of prymnesiophyte sequences and both are always higher than the fraction of prymnesiophyte biovolume concentrations. Fuco/∑ *pigs* ratios, relative diatom sequence abundance, and relative diatom biovolume concentrations are similar across samples, as are 19ButFuco/∑ *pigs* ratios and relative dictyochophyte + pelagophyte sequence abundance. Cryptophytes are consistently a small fraction of PCC from all three methods, with the exception of a few samples during NAAMES exhibiting higher relative cryptophyte biovolume concentrations (Figure 2C). Finally, at NAAMES3 stations 1 and 2, there is a notable maximum in the relative fraction of Zea/∑ *pigs* (a picophytoplankton and cyanobacteria marker pigment; Figure 2A), which could not be revealed by the other 2 methods, as this group is not quantified by the IFCB or 18S methods.

The above noted qualitative comparisons of absolute values (except 18S amplicons) across the dataset (Figure 1) and relative values between samples (Figure 2) demonstrate broad similarities and notable differences among the three methods. However, these comparisons consider only the component of the dataset that includes known information relating to phytoplankton groups: the methods also measure other pigments (Figure 1A, 2A), sequences (Figure 1B, 2B), and imaged cells (Figure 1C, 2C). The “other” accessory pigments from HPLC (Chlc12, Chlc3, DVchla, DVchlb, Neo, Viola, Pras; most of which are expressed by the taxa already resolved by other biomarker pigments) are a consistent fraction of the total pigment concentration (32-48%; mean = 37%; median = 36%). Similarly, the “other” sequences from 18S (including unclassified sequences) are a small fraction of the total sequence abundance (2-18%; mean = 9%; median = 8%). Conversely, the “other” cells from IFCB compose a sometimes large fraction of the total IFCB biovolume concentration including unclassified or unidentified images (20-83%; mean = 45%; median = 41%) that covaries with the total IFCB biovolume concentration for a given sample (R^2^ = 0.90).

### 3.2 Covariation of PCC from pigments, 18S amplicons, and IFCB group biovolume concentrations

Relationships between relative pigment ratios and relative sequence abundances are significant (p<<0.001), positive, and strong for diatoms (Figure S2A; Table 3; R^2^ = 0.57), dictyochophytes + pelagophytes (Figure S2C; R^2^ = 0.60), and chlorophytes (Figure S2E; R^2^ = 0.59). Relationships are also significant (p<0.001) and positive for prymnesiophytes (Figure S2D; R^2^ = 0.37) and cryptophytes (Figure S2F; R^2^ = 0.41). Dinoflagellates have the weakest positive relationship of the groups considered here (Figure S2B; R^2^ = 0.13; p = 0.01).

**Table 3.**
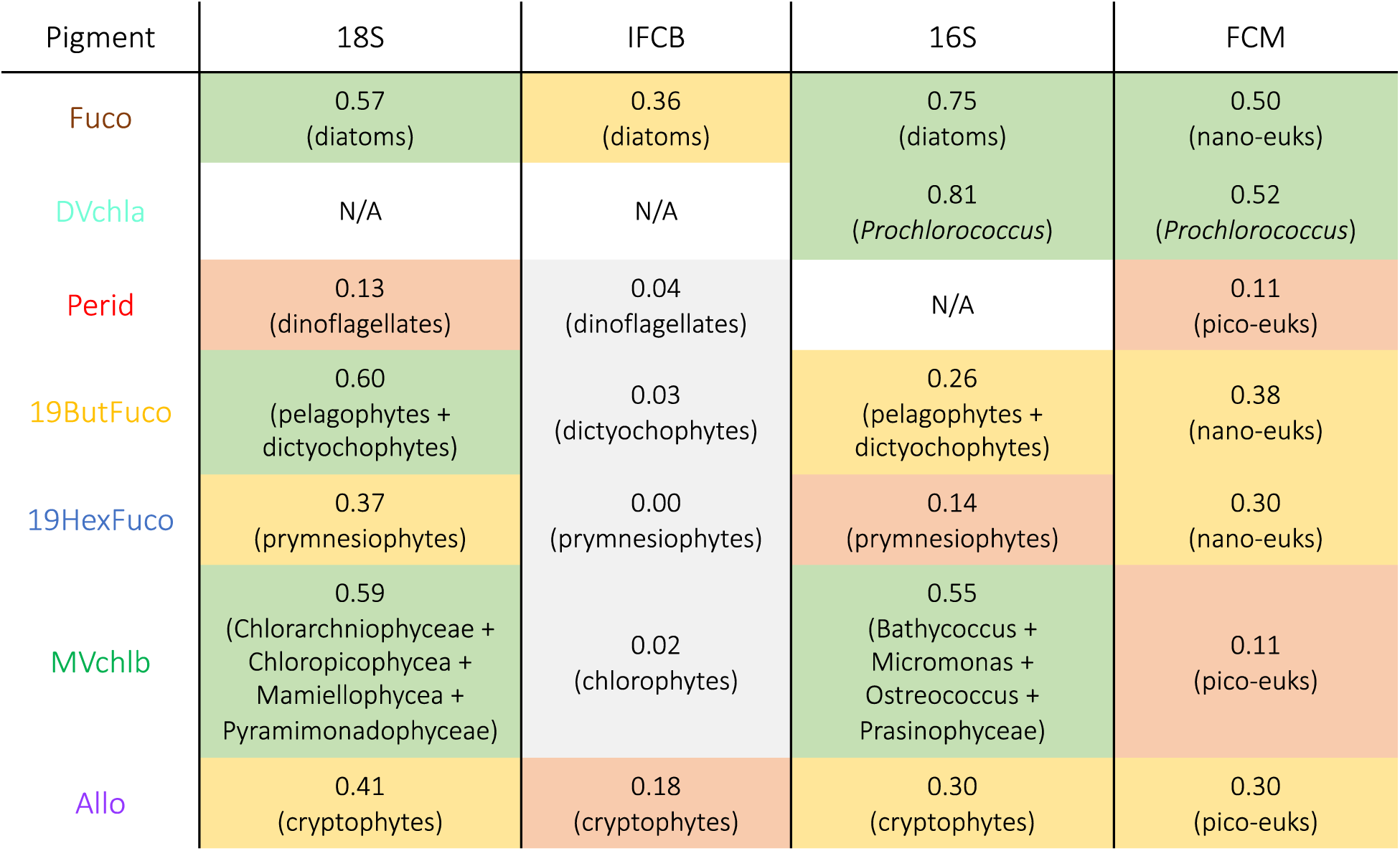
Pearson’s correlation coefficient (R^2^) between ratios of pigments to Tchla and other PCC methods (18S, IFCB, 16S, and FCM). Shades indicate to the relative strength of the relationship: green for R^2^ > 0.5, yellow for 0.5 > R^2^ > 0.25, red for 0.25 > R^2^ > 0.10, and grey for R^2^ < 0.10. N/A indicates that the PCC method does not have a corresponding measurement for that pigment.

| Pigment | 18S | IFCB | 16S | FCM |
| --- | --- | --- | --- | --- |
| Fuco | 0.57<br>(diatoms) | 0.36<br>(diatoms) | 0.75<br>(diatoms) | 0.50<br>(nano-euks) |
| DVchl<sub>a</sub> | N/A | N/A | 0.81<br>(*Prochlorococcus*) | 0.52<br>(*Prochlorococcus*) |
| Perid | 0.13<br>(dinoflagellates) | 0.04<br>(dinoflagellates) | N/A | 0.11<br>(pico-euks) |
| 19ButFuco | 0.60<br>(pelagophytes +<br>dictyochophytes) | 0.03<br>(dictyochophytes) | 0.26<br>(pelagophytes +<br>dictyochophytes) | 0.38<br>(nano-euks) |
| 19HexFuco | 0.37<br>(prymnesiophytes) | 0.00<br>(prymnesiophytes) | 0.14<br>(prymnesiophytes) | 0.30<br>(nano-euks) |
| MVchl<sub>b</sub> | 0.59<br>(Chlorarchniophyceae +<br>Chloropicophyceae +<br>Mamiellophyceae +<br>Pyramimonadophyceae) | 0.02<br>(chlorophytes) | 0.55<br>(Bathycoccus +<br>Micromonas +<br>Ostreococcus +<br>Prasinophyceae) | 0.11<br>(pico-euks) |
| Allo | 0.41<br>(cryptophytes) | 0.18<br>(cryptophytes) | 0.30<br>(cryptophytes) | 0.30<br>(pico-euks) |

Qualitatively, there are some similarities between the relative IFCB biovolume concentrations and the relationships between relative pigment concentrations and relative sequence abundances. For instance, in most cases, the highest relative diatom biovolume concentrations correspond to the highest Fuco/Tchla concentrations and largest relative diatom sequence abundance (Figure S2A). However, statistical relationships between relative pigment concentrations and biovolume concentrations for these same groups (Figure S3) are either weak (for diatoms and cryptophytes) or statistically insignificant (for all other groups).

A chord diagram (Gu et al., 2014) demonstrates the relative strength of the correlations of pigment ratios with class-level relative sequence abundances and relative IFCB biovolume concentrations (Figure 3). The “other” fraction of the IFCB is also included, to consider relationships between pigments and unidentifiable cells. The width of the edge between each pigment and 18S class or IFCB group describes the relative strength of the correlation between those groups. Many biomarker pigments share edges with the class or group that they are expected to represent. For instance, Fuco is strongly associated with relative diatom sequence abundance and IFCB diatom biovolume concentration. Allo is associated with relative cryptophyte sequence abundance and IFCB cryptophyte biovolume concentration. 19ButFuco shares edges with relative pelagophyte and dictyochophyte sequence abundances, while 19HexFuco shares edges with relative prymnesiophyte sequence abundance. MVchlb and other chlorophyte accessory pigments (Neo, Viola, Pras) share edges with most MVchlb-containing classes (Chloropicophyceae, Chorarachniophyceae, and Mamiellophyceae), as well as with relative IFCB chlorophyte biovolume concentrations.

**Figure 3.**
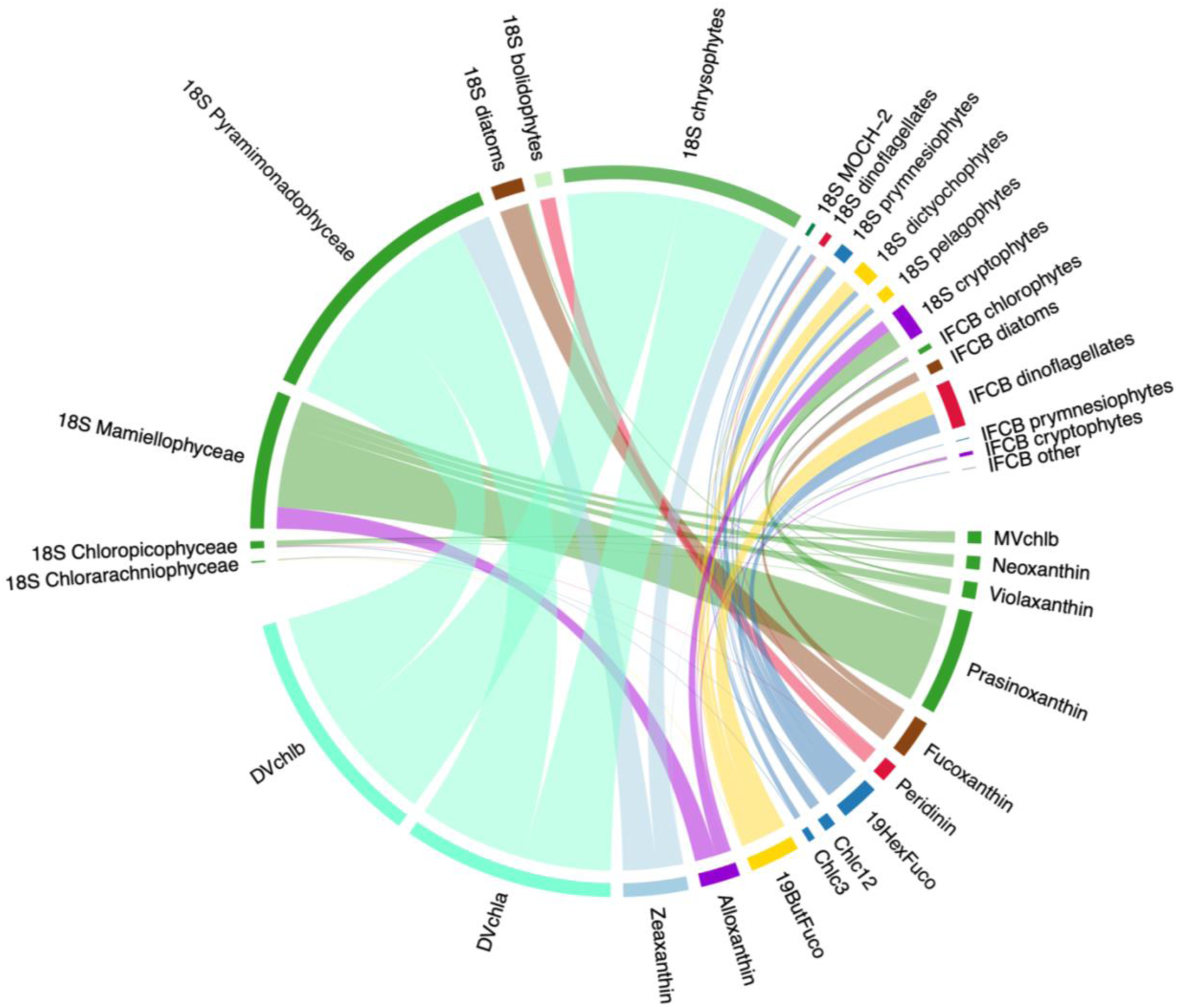
Chord diagram constructed from the weighted adjacency matrix of HPLC pigments (normalized to Tchla), class-level 18S amplicon sequencing (relative sequence abundances), and IFCB groups (relative biovolume concentration) from NAAMES and EXPORTS. The diagram is directed from pigments to other methods; line colors correspond with pigments. The width of the line connecting pigments to 18S classes or IFCB groups is based on the weighted correlation coefficient among these parameters (max = 0.87 from DVchlb to Pyramimonadophyceae; min = 0.006 from Perid to Chloropicophyceae). Label colors are consistent with Figure 1.

The chord diagram (Figure 3) also reveals significant covariation between taxa, often with unexpected associations among pigments and higher-resolution PCC methods. For instance, the picoplankton biomarker pigments (Zea, DVchla, DVchlb) are unexpectedly associated with one green algal class (Pyramimonadophyceae) and with chrysophyte relative sequence abundance. Similarly, Perid is strongly associated with bolidophytes, which are pico-phytoplankton known to contain Fuco but not Perid and thus more often associated with diatom biomarkers (Kuwata et al., 2018), though not in this dataset. 19ButFuco and 19HexFuco are both associated with the relative IFCB dinoflagellate biovolume concentration, though dinoflagellates are not known to contain either of these pigments unless acquired through mixotrophy (e.g., Nascimento et al., 2005). Finally, MOCH-2 (a red algal class) and IFCB “other” biovolume concentrations both share an edge with 19HexFuco.

The information contained in the chord diagram can be further visualized with an unweighted graph that considers the strongest connections among variables and across methods while still prioritizing the strongest within-group connections (Figure 4). This unweighted graph separates pigment ratios, relative 18S sequence abundances, and relative IFCB biovolume concentrations by highlighting positive connections between groups and demonstrating relative distances between broad communities. Six communities separate on the basis of network-based community detection analysis. The first community (brown diamonds in Figure 3) includes Fuco, diatom sequence abundance, and IFCB diatoms. The second community (light blue circles) is made up of cyanobacterial pigments (Zea, DVchla, DVchlb) and two 18S classes: Pyramimonadophyceae (a green algal class) and chrysophytes (a red algal class). This association in the second (light blue circles) community is not surprising given the consistently strong correlations among these variables across analyses (Figure 3). The third community (light green triangles) is mostly composed of pigments and 18S classes in the cryptophyte and green algal groups: Allo, 18S cryptophytes, and IFCB cryptophytes; MVchlb, Neo, Viola, Pras, Mamiellophyceae, Chloropicophyceae, and IFCB chlorophytes. This third community also unexpectedly includes IFCB dictyochophytes, but this group is also arranged closely, and more expectedly, to the fourth community (dark blue squares), which includes dictyochophytes + pelagophytes, prymnesiophytes, and some dinoflagellate markers. The fourth community (dark blue squares) comprises: 19HexFuco, Chlc12, Chlc3, and 18S prymnesiophytes; 19ButFuco, 18S dictyochophytes, and 18S pelagophytes; and 18S dinoflagellates and IFCB dinoflagellates.

**Figure 4.**
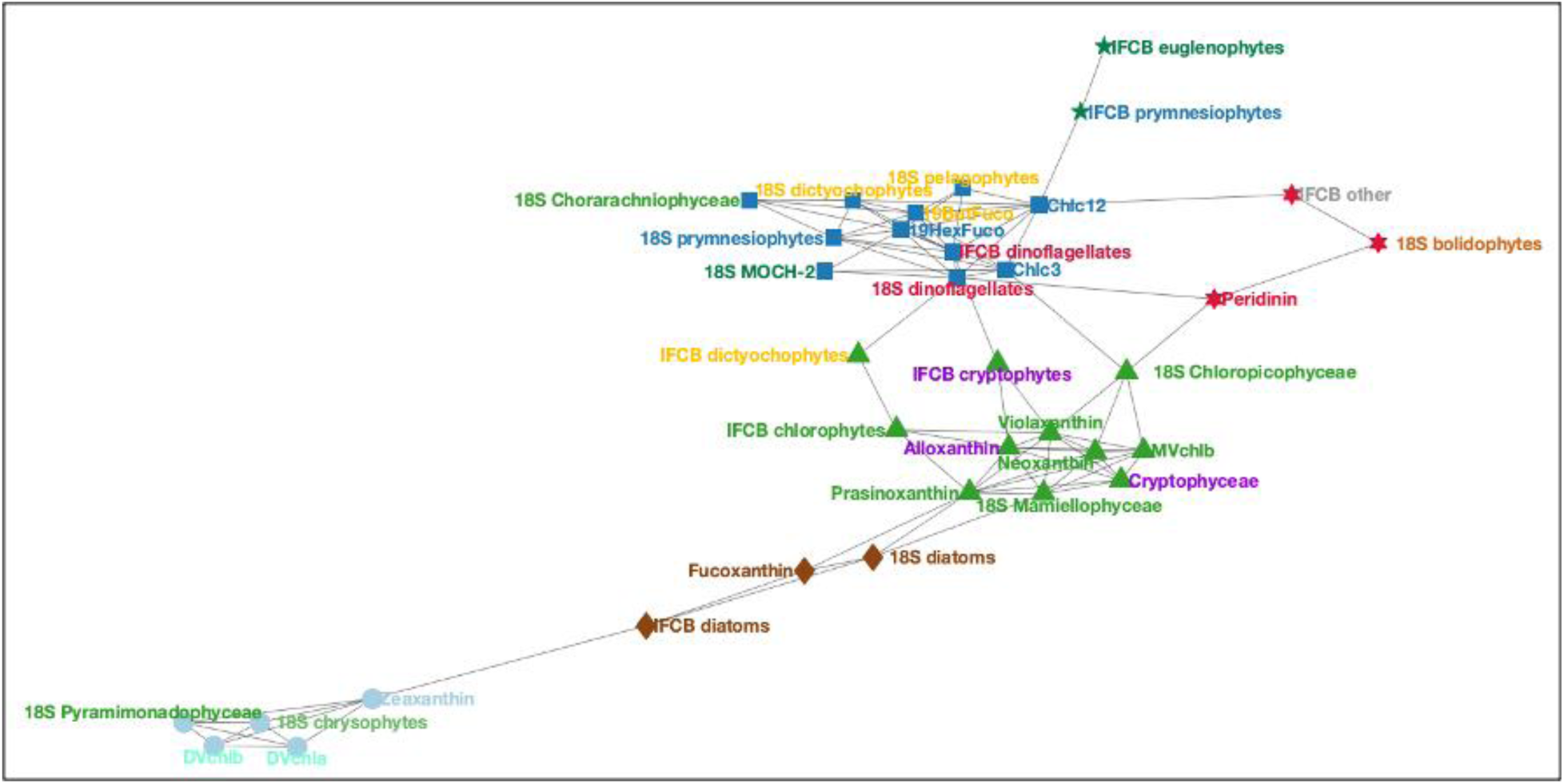
Unweighted graph built from the adjacency matrix of HPLC pigments (normalized to Tchla), 18S (relative sequence abundances), and IFCB (relative biovolume concentration) from NAAMES and EXPORTS. Node colors and shapes are determined by the community assignment from network-based community detection analysis. Label colors are consistent with Figure 1.

MOCH-2 and Chlorarachniophyceae are also associated with this community, which is expected given the correlations between these 18S classes and 19HexFuco in other statistical analyses (Figure 3). The fifth community (dark green five-point stars) contains IFCB prymnesiophytes and IFCB euglenoids. These two latter classes are relatively sparse within the dataset and cluster closely across analyses. Finally, the sixth community (red six-point stars) is composed of IFCB “other,” along with Perid and 18S bolidophytes, mirroring a surprising association between the latter two groups found in the chord diagram. IFCB “other” is also connected to Chlc12 in the dark blue squares community.

### 3.3 Trends in PCC from HPLC pigments, 16S amplicons, and FCM cell counts

A similar comparison was performed for the dataset made up of HPLC pigments, 16S amplicon sequencing, and FCM from the NAAMES cruises showing a mix of good and poor correspondence across methods. Median relative abundances of *Prochlorococcus* sp. are similar across all three methods (Figure 5A, C, E). However, the relative fraction of DVchla is often lower than the relative sequence abundance or cell counts of *Prochlorococcus* from the other two methods (Figure 5B, D, F). There are also similar median fractions of Zea, *Synechococcus* sp. from 16S, and *Synechococcus* sp. from FCM, though Zea is not unique to *Synechococcus*. In some samples (e.g., early transit on NAAMES 4; see Figure 5F axis labels), the relative Zea concentration is much higher than the fraction of *Synechococcus* from 16S or FCM. In other samples (e.g., mid-cruise transit during NAAMES 4; see Figure 5F axis labels), the opposite trend is observed. There are similar contributions to PCC by green algae, diatom, prymnesiophyte, and dictyochophyte + pelagophyte markers between pigments and 16S (Figure 5A, C, E). However, the relative fractions of these groups across individual samples are often quite different. For example, the relatively low fraction of prymnesiophyte sequences compared to the relatively high fraction of 19HexFuco to other accessory pigments is particularly notable (Figure 5B, D).

**Figure 5.**
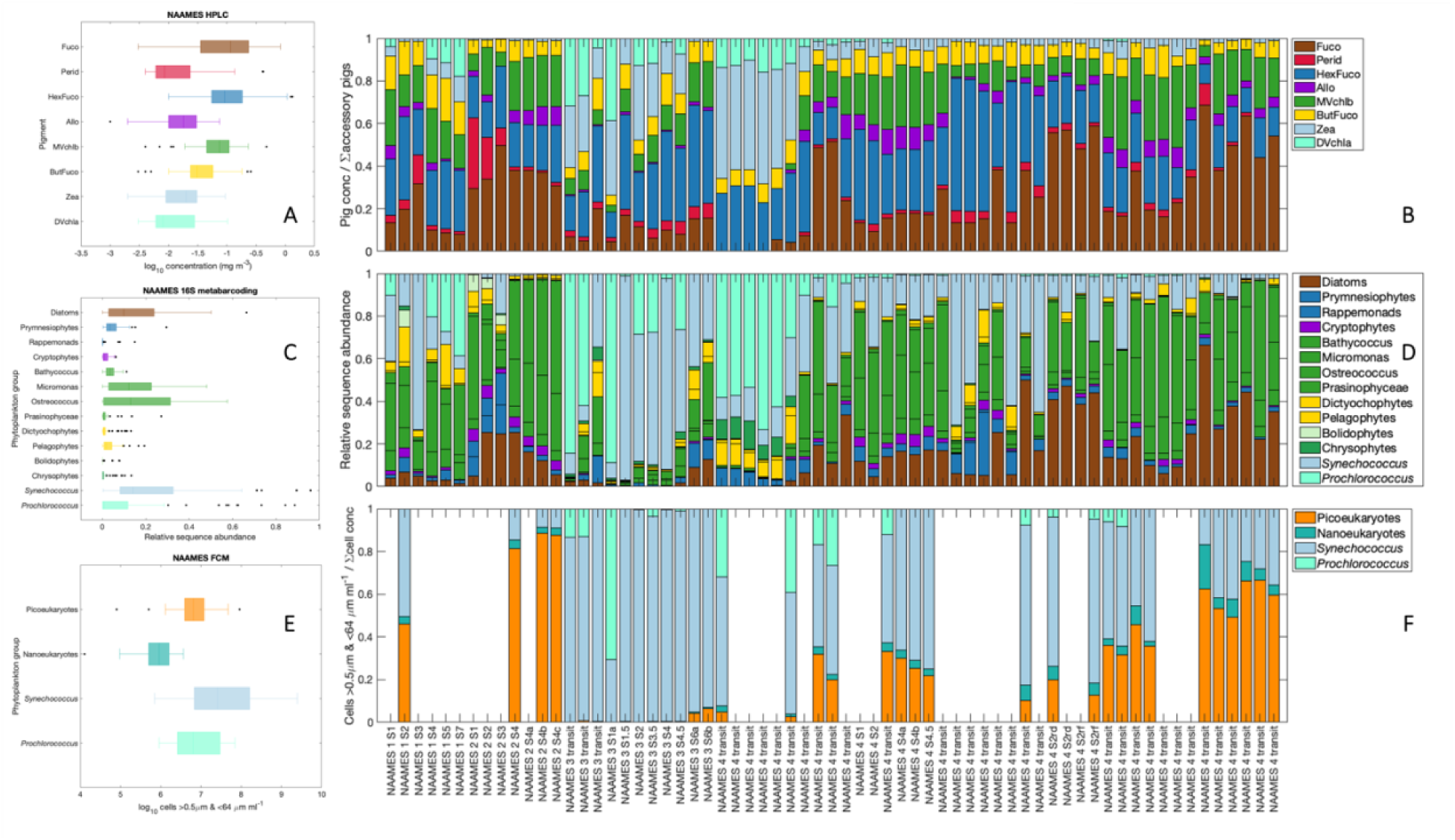
(A) Concentrations and (B) relative fractions of phytoplankton pigments; (C) and (D) relative sequence abundances from 16S; and (E) cell counts and (F) relative fractions of cells measured by FCM, all from NAAMES. Samples are organized from left to right in the order collected, from NAAMES 2 and 3 on the left half and NAAMES 4 on the right half. Boxes and fractions are colored similarly for shared groups: diatoms in brown, prymnesiophytes in dark blue, cryptophytes in purple, chlorophytes in bright green, dictyochophytes + pelagophytes in gold, *Synechococcus* in light blue, and *Prochlorococcus* in cyan.

### 3.4 Covariation of PCC from pigments, 16S amplicons, and FCM cell counts

As with the HPLC, 18S, and IFCB dataset, the qualitative comparisons among HPLC pigment ratios, 16S relative sequence abundances, and FCM cell count fractions show broad patterns of agreement among groups and across methods (see Table 1 for assumed group assignments with each method). Pigment concentrations were normalized to their sum here (which is highly correlated with Tchla in this dataset; R^2^ = 0.93) for comparison with the other two methods. The direct quantitative comparison between pigment-based PCC and 16S amplicon sequencing reveals significant relationships (p<<0.001) for some groups (Table 3; Figure S4). Diatoms (Figure S4A; R^2^ = 0.75), green algae (Figure S4E; R^2^ = 0.57), and *Prochlorococcus* (Figure S4F; R^2^ = 0.81) are highly positively correlated across methods. Correlations between pigment and 16S relative PCC contributions for cryptophytes (Figure S4C; R^2^ = 0.30) and dictyochophytes + pelagophytes (Figure S4D; R^2^ = 0.26) are statistically significant (p<0.001) and positive, but with poorer linear regression fit statistics. Besides dinoflagellates, the weakest positive relationship of the groups considered here is found for prymnesiophytes (Figure S4B; R^2^ = 0.14; p = 0.002). There are also strong positive relationships between Fuco/Tchla and nanoeukaryote cell fractions from FCM (R^2^ = 0.50) and between DVchla/Tchla and *Prochlorococcus* from FCM (R^2^ = 0.52). To a lesser degree, Allo/Tchla and picoeukaryote cell fractions from FCM are also positively correlated (R^2^ = 0.30). Zea/Tchla and *Synechococcus* are only weakly correlated (R^2^ = 0.10) and there are no other notable correlations between flow cytometry cell fractions and pigment-based PCC (Tables 1-2).

A chord diagram was constructed to show the relative strength of the weighted correlations among pigment-based PCC and PCC from 16S and FCM (Figure 6). Many of the connections in this diagram are expected based on the distribution of pigments in major phytoplankton groups. *Prochlorococcus* from 16S and from FCM are strongly correlated with DVchla, DVChlb, and Zea. As expected, we also find that Fuco shares an edge with 16S diatoms, 19HexFuco shares an edge with 16S prymnesiophytes, 19ButFuco shares an edge with 16S pelagophytes, and Allo shares an edge with 16S cryptophytes. All four green algal pigments are correlated with the chlorophyte classes from 16S. There are also unexpected correlations between groups (Figure 6). For instance, Zea is strongly correlated with 16S chrysophytes (as in the HPLC and 18S dataset; Figure 3) and with 16S dictyochophytes. Likewise, Perid shares edges with 16S bolidophytes (as in the HPLC and 18S dataset; Figure 2) and with rappemonads (a red algal class that contains Fuco, 19HexFuco, and chlorophyll c; Kawachi et al., 2021), but we have found no evidence in the literature that members of these classes contain Perid.

**Figure 6.**
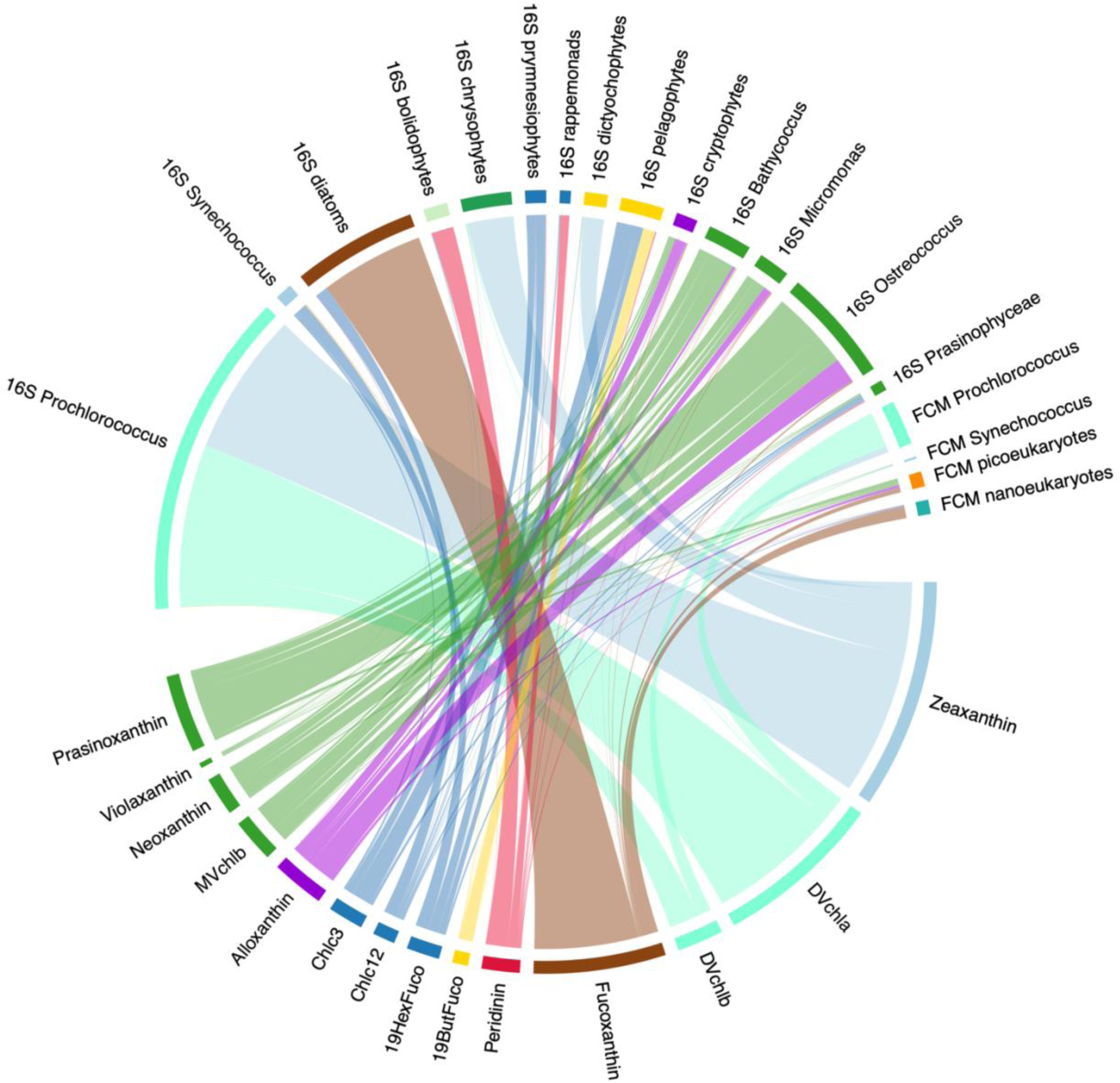
Chord diagram constructed from weighted adjacency matrix of HPLC pigments (normalized to Tchla), group level 16S (relative sequence abundances), and FCM (relative fraction of cells) from NAAMES. The diagram is directed from pigments to other methods; line colors correspond to pigments. The width of the line connecting pigments to 16S or FCM groups is based on the weighted correlation coefficient among these parameters (max = 0.59 from Zea to *Prochlorococcus*; min = 0.007 from 19HexFuco to Prasinophyceae). Label colors are consistent with Figure 4.

Interestingly, we find that *Synechococcus* from 16S is correlated with 19HexFuco, Chlc12, and Chlc3, while *Synechococcus* from FCM is correlated with Zea, as expected. The picoeukaryote fraction of the FCM dataset shares edges with green algal pigments, Allo, and Fuco, while the nanoeukaryote fraction shares edges with Allo, prymnesiophyte pigments, and Fuco.

As a final analysis, a graph was constructed to visualize the relative correlations among communities of pigments, 16S groups, and FCM groups (Figure 7). Five broad communities separated from a network-based community detection analysis. The first community (cyan circles) comprises cyanobacterial markers: Zea, DVchla, DVchlb, and *Prochlorococcus* from 16S and from FCM. This community also includes 16S chrysophytes and dictyochophytes, presumably due to their strong correlations with Zea (Figure 6). The second community (green triangles) is composed of chlorophyte and cryptophyte pigments and 16S groups: Allo and cryptophytes; MVchlb, Neo, Viola, Pras, *Micromonas* spp., *Bathycoccus* spp., and *Ostreococcus* spp. This second community is highly connected to picoeukaryotes, which belong to the third community (brown diamonds) along with nanoeukaryotes and diatom pigments (Fuco, Chlc12) and 16S diatoms. Chlc12 links the brown diamonds community to the dark blue squares community, which includes prymnesiophyte, dictyochophyte, and pelagophyte pigments and 16S groups (19HexFuco, 19ButFuco, Chlc3, prymnesiophytes, pelagophytes). The dark blue squares community also includes Prasinophyceae (a chlorophyte class) and *Synechococcus* from 16S and FCM. Finally, the sixth community (red six-point stars) includes Perid, 16S bolidophytes, and 16S rappemonads, similarly to the chord analyses (Figure 6).

**Figure 7.**
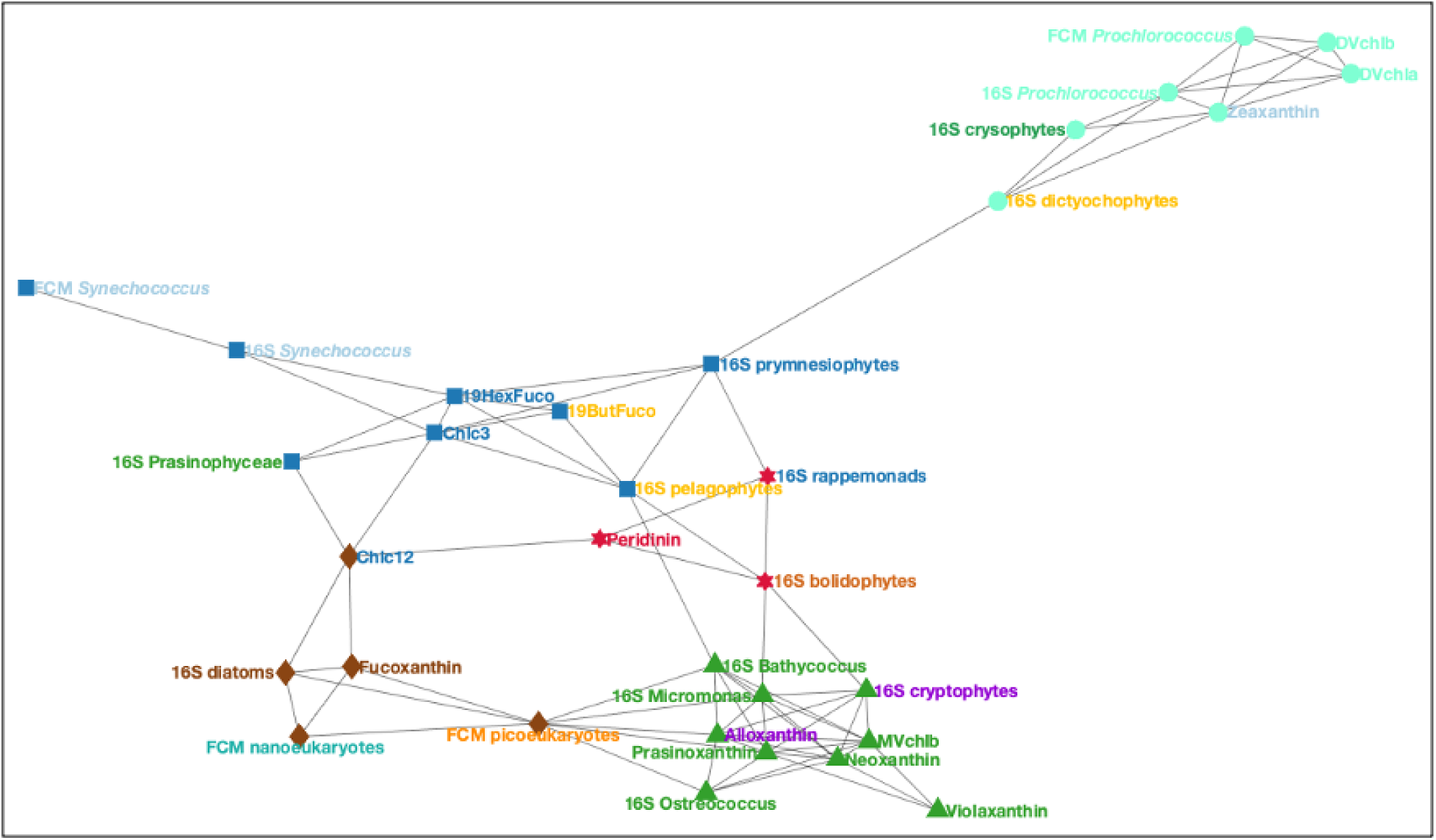
Unweighted graph from adjacency matrix of HPLC pigments (normalized to Tchla), 16S (relative sequence abundances), and FCM (relative cell counts). Node colors and shapes are set by the community assignment from network-based community detection analysis. Label colors are consistent with Figure 4.

## 4. Discussion

### 4.1 Overview

The goal of this study was to assess the correspondence of pigment-based PCC with PCC metrics determined by other methods, often with higher taxonomic resolution. Taken together, these analyses reveal broad correspondence among pigment-based PCC and other methods at the class-to group-level for most cases examined (Table 3; Figure S2; Figure S4). The ratio of the expected biomarker pigments to Tchla was well correlated with the relative sequence abundance of the associated class, with the notable exceptions of dinoflagellates from 18S and prymnesiophytes from 16S. Strong positive correlations were found between pigment ratios and relative IFCB biovolume concentrations for diatoms (Figure S3) and between relative pigment concentrations and fractions of FCM cell counts for *Prochlorococcus*. While these results reveal many of the expected correlations among accessory pigments and other PCC methods, we also observed unexpected correlations between other pigments and phytoplankton groups.

In the sections that follow, we use examples from the current datasets to investigate some sources of inconsistencies among methods that may be associated with uncertainty in pigment-based PCC analyses. Inconsistencies among methods can provide opportunities to further quantify the challenges of pigments as biomarkers for specific phytoplankton groups (e.g., the outliers of the Perid vs. 18S dinoflagellates relationship; Figure S2B) or to describe the co-occurrence of some groups in their environment (e.g., the associations of DVchla and DVchlb with some 18S classes; Figures 2A, 4-5). We discuss the major strengths and weaknesses for each method highlighted by this analysis and provide some recommendations for PCC method selection in particular use cases. Finally, we review the challenges and impediments to integrating PCC methods, particularly for calibration and validation of ocean color models, for which HPLC pigments remain the most common approach. Of the methods considered here, no single approach provides “perfect” PCC assessment. The hope is that when methods are combined, however, a more robust characterization of PCC can be achieved.

### 4.2 Comments on challenges with pigment-based PCC determinations

There are various reasons why different PCC methods can produce disparate results. Here, we summarize four possible sources of inconsistencies between pigment-based PCC and other PCC determinations. We use correlation analysis of ASVs with environmental parameters and pigments to demonstrate the complexities in interpreting PCC dynamics due to these sources of inconsistency.

First, intra-group variations in phytoplankton pigment composition and concentration arise when different phytoplankton species from the same class express different suites or amounts of accessory pigments (e.g., Zapata et al., 2004; Irigoien et al., 2004; Zapata et al., 2012; Neeley et al., 2022). While there might be broad agreement between pigments and relative sequence abundances or relative biovolume concentrations at the class level, many of these relationships change or vanish at the genus-to species-level, making biomarker pigments a limited taxonomic resource. Secondly, there are inter-group variations in phytoplankton pigment composition and concentration (Jeffrey et al., 2011 and references therein) since many groups share accessory pigments. For example, Fuco is often used as a biomarker for diatoms but is also found in prymnesiophytes, chrysophytes, and pelagophytes, as well as some dinoflagellates, dictyochophytes, pelagophytes, and bolidophytes. Feeding strategies such as mixotrophy, through which a phytoplankter might acquire pigments that are not typically found in that group (Stoecker et al., 2017; Li et al., 2022), can also drive inter-group variation in pigment composition. Third, some genera or species may co-occur in the environment, leading to the covariation of unexpected taxa with a pigment that they may not contain, but others in the group do. Finally, phytoplankton pigments may vary in composition and concentration due to phytoplankton physiological responses to the physical environment (Thompson et al., 2007), which includes light history (particularly as many pigments have photoprotective functions, including Allo and Zea) and nutrient availability (Schlüter et al., 2000; Henriksen et al., 2002; Catlett et al., 2022).

To explore intra-group variations in pigment expression, we examine correlations of individual ASVs with pigments in the 18S dataset (Figure 8). This approach contrasts with the analysis of aggregate class-or group-level taxonomy (as shown in Figures 3-4). Here, we compare the relative abundance of the 135 ASVs that each comprise >1% of the total sequences in any given sample in this dataset with pigment ratios to Tchla. While there are broad patterns that mirror the positive class-level correlations between pigments and relative sequence abundances, correlations are highly variable within classes. For instance, about half of the prymnesiophye ASVs are positively correlated with 19HexFuco (including 8 of the 10 most abundant ASVs), while the other half are negatively correlated (Figure 8D). Similarly, despite the strong relationship between Fuco/Tchla and relative diatom sequence abundance (Table 3; Figure S2A), there are many diatom ASVs that are insignificantly or weakly negatively correlated with Fuco. The variability in these relationships highlights the difficulty of comparing relative abundances in correlation space. Relative abundances of one ASV are necessarily dependent on the rest of the community in the sample or dataset. This analysis used all ASVs that were >1% abundant in the dataset, meaning that some ASVs were only present in a small fraction of the samples (Figure 8B) or only ever reached a very low overall abundance in the dataset (Figure 8C). The dominant ASVs drive the relationship at the group level between pigments and ASVs, but not all ASVs within a group will be correlated with the expected biomarker pigment. Thus, it is perhaps unsurprising that the correlations between pigments and relative sequence abundances are variable across all ASVs, as the relative abundances themselves are highly variable.

**Figure 8.**
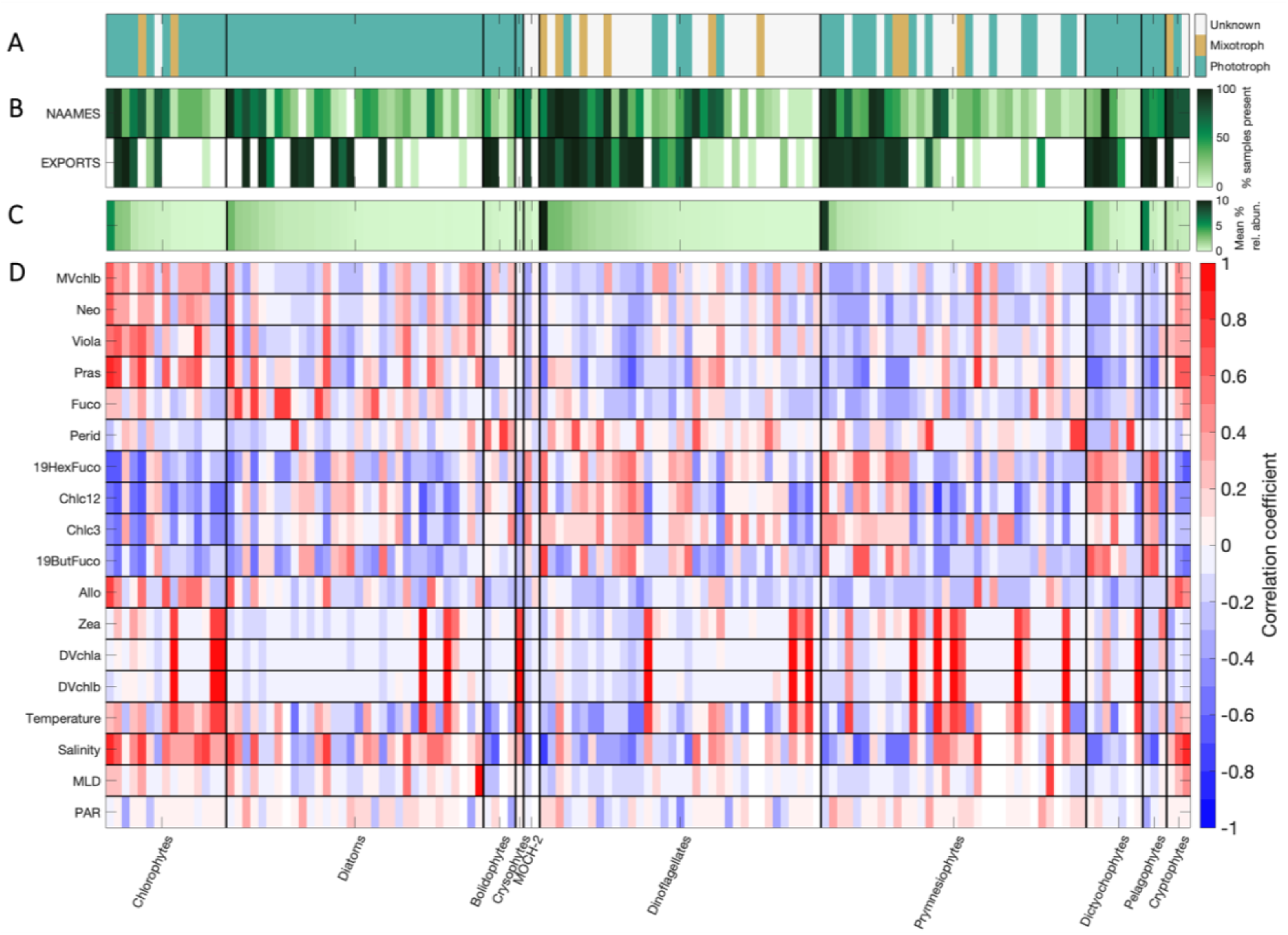
(A) Presumed feeding strategy for each >1% abundant ASV (teal = known phototroph, orange = known mixotroph, white = unknown). (B) The relative frequency of each ASV on NAAMES vs. EXPORTS. (C) Mean relative percent abundance of each ASV in the dataset. (D) Pearson’s correlation coefficient (R) between relative pigment concentrations and ASVs from 18S (relative sequence abundances, sorted by mean abundance within each class). The strength of the correlation is shown on a scale from −1 (blue) to 1 (red). Correlations with environmental variables (temperature, salinity, MLD, PAR) are also shown.

Many of the ASVs in the 18S dataset were classified as mixotrophs (Figure 8A) or have undocumented feeding strategies but are members of groups with mixotrophic representatives. Members of many of the classes represented in this dataset have demonstrated mixotrophy in nature or in culture. For instance, a recent study demonstrated the phagocytosis of *Prochlorococcus* sp. by dictyochophytes, prymnesiophytes, chlorophytes, chrysophytes, bolidophytes, and dinoflagellates (Li et al., 2022). Some of the ASVs in these classes have strong correlations with DVchla and DVchlb, which are marker pigments for *Prochlorococcus* (Figure 8D). Of the sixteen ASVs that are highly correlated with DVchla and DVchlb (R>0.7), eight were classified as phototrophs, two as mixotrophs (a chlorophyte, *Cymbomonas tetramitiformis*, and a prymnesiophyte, *Chrysochromulina acantha*), and six have undocumented feeding strategies but are members of groups known to include mixotrophs (specifically, three dinoflagellate ASVs and three prymnesiophyte ASVs). This dataset only indicates correlations between these ASVs and pigments and there may be other reasons for this correspondence, but mixotrophic assimilation of *Prochlorococcus* pigments is one possibility.

There is also the possibility of environmental co-occurrence between taxa. Some unlikely correspondences between pigments and other PCC determinations may arise from a near-random sampling of an evolving PCC distribution. PCC can be highly variable in space and time. Each sample represents a snapshot of the specific environment at one moment in time and one position in space, and thus can be limited in its ability to capture the broader context (e.g., Siegel et al., 2001; Estapa et al., 2015). The correlative analyses applied here identify statistical relationships between pigments and taxa that co-occur, not necessarily within a taxonomic group but within a covarying community. The associations, for instance, of Perid with bolidophytes from both 18S and 16S or Zea with chrysophytes from both 18S and 16S (Figures 4 and 7) are not attributable to any documented pigment-based taxonomy and thus possibly reflects environmental covariation in these analyses that leads to a high correlation among these parameters.

Environmental data can also provide further insights into relationships between PCC methods. *Prochlorococcus* relative sequence abundance is highly positively correlated with its marker pigments DVchla and DVchlb, but also with sea surface temperature (Figure S5). Many of the 18S ASVs that have strong positive correlations with DVchla and DVchlb (but are not expected to contain these pigments) are also positively correlated with sea surface temperature (Figure 8D), suggesting a co-occurrence of these 18S ASVs with *Prochlorococcus* in the environment, as evidenced by the biomarker pigments and the warm ocean temperature. In this anecdote, the combined PCC methods validate the pigment-based PCC, but also draw upon environmental co-variability to inform a more complete picture of PCC. These relationships among disparate parameters are also useful for considering these datasets in the context of community ecology, where interactions between phytoplankton shape the ecosystem as a whole (e.g., Lima-Mendez et al., 2015; Zhou and Ning, 2017). However, correlations between pigments and other methods are unable to separate the mechanism of covariation (e.g., whether it represents a common response to the environment or a biological interaction).

The oceanographic context from which the samples were collected can also inform associations between taxonomic groups. In this study, mixed layer depth (MLD) and PAR were typically weakly correlated with individual 18S ASVs (Figure 8D), though relative diatom sequences from 16S were positively correlated with MLD and PAR (Figure S5), as were some chlorophyte classes. As the PCC methods compared here included cell-specific measurements from the IFCB and FCM, the impact of environmental conditions could be indirectly interrogated by examining changes in pigment-per-cell or pigment-per-biovolume concentration over the dataset that might be associated with variability in the light environment. For instance, when the outliers from the Perid/Tchla vs. relative dinoflagellate sequence abundance relationship (Figure S2B; highest outlier circled in red) are considered as a function of pigment-per-biovolume, there is anomalously high Perid-per-biovolume in those samples (Figure S6A), while the Tchla-per-biovolume for the outlier samples is consistent with the mean value for the dataset (Figure S6B). Together, this result suggests that these samples comprise Perid-containing dinoflagellates with higher Perid per cell than the rest of the dataset. This trend in the outlier samples may also be due to intra-group variability in pigment concentration or to responses of dinoflagellate pigmentation to environmental stimuli, with some dinoflagellate ASVs in those outlier samples containing higher ratios of Perid/Tchla than the mean in the dataset. The most abundant ASVs in this sample include two dinoflagellates (*Biechelaria* sp. and *Prorocentrum* sp.), but we could not find evidence in the literature to support these genera having considerably larger Perid/Tchla ratios than other Perid-containing phototrophic dinoflagellates.

Ultimately, inconsistencies in the correlations between pigments and other PCC methods may have arisen as a combination of factors in the environment and ecosystem (due to physical mixing, mixotrophy, co-occurrence with other groups, etc.) that cannot be easily disentangled in the present dataset. It is also important to note that correlation-based analyses may be poorly suited for compositional datasets, such as the ones used here, and spurious correlations may confuse the interpretation of the results for ecological data (Gloor et al., 2017; Hunter-Cevera et al., 2021). Finally, each method has errors and uncertainties associated with sample collection, processing, analysis, and taxonomic assignment which introduce uncertainties into the PCC metrics as defined and thus will impact the associations between groups. A complete propagation of all possible sources of error for each method would potentially impact the correlations and comparisons between observations and methods.

### 4.3 Recommendations for measuring PCC

The results of this analysis suggest that the most complete picture of PCC will be achieved with a combination of methods, particularly given the complementary strengths and limitations of the common methods for describing and quantifying phytoplankton communities. Thus, the choice of method(s) to be used for a given application is a function of the desired taxonomic and spatio-temporal resolution, the purpose of the study, the cost of the analysis, and the time scale for analytical results. For instance, the IFCB has been used successfully to detect (Campbell et al., 2010) and monitor the development (Brosnahan et al., 2015) of harmful algal blooms (HABs; e.g., Anderson et al., 2012) in varying ecosystems. IFCB data are available in near real-time, which allows for quick detection and timely warnings when a harmful bloom develops (as opposed to methods such as pigments or amplicon sequencing, which require weeks to months of processing and analysis after sample collection). Alternatively, at a time-series observatory where the goal is long-term monitoring of the seasonal succession of phytoplankton and changes in PCC over time with environmental change, a combination of methods could be appropriate. Pigments allow for comparison with ongoing optical measurements alongside a record of PCC (Zhang et al., 2015; Catlett et al., 2021b), while amplicon sequencing focuses on high taxonomic resolution at the site (Needham and Fuhrman, 2016; Yeh and Fuhrman, 2022).

Continuous flow-through phytoplankton imaging systems, such as IFCB, could also be used in combination with other methods, such as FCM, to acquire high-resolution PCC across a large range of cell sizes with samples collected approximately every twenty minutes (e.g., Peacock et al., 2014; Hunter-Cevera et al., 2016). Sometimes, the impact of an environmental disturbance on the phytoplankton community may be the focus of an investigation. In these cases, a combination of real-time imaging approaches, amplicon sequencing, and/or pigment data can confirm the impact of a disturbance on the function or optical properties of the phytoplankton community (e.g., Laney and Sosik, 2014; Kramer et al., 2020b). Across timescales, a combination of methods can offer a more nuanced picture of PCC.

Standardized PCC information is essential for models of carbon export or the biological pump, which typically include phytoplankton size and/or community composition terms to constrain the export of phytoplankton carbon from the surface ocean to the deep ocean (e.g., Guidi et al., 2015; Durkin et al., 2022; Siegel et al., 2023). Many Earth system models use satellite data to achieve global ocean coverage, and pigment concentrations are currently the only PCC metric derived from ocean color data (Kramer et al., 2022). Thus, pigment measurements remain important to link ocean color estimates of PCC to in-water data. Methods that directly measure cell biovolume concentration (IFCB, FCM) are also useful to estimate carbon-per-cell (Menden-Deuer and Lessard, 2000). Direct comparisons between satellite remote sensing observations, pigment concentrations, and higher-resolution PCC methods are rare, but will be essential to constrain model outputs (Chase et al., 2022), particularly as sequencing methods improve and become more quantitative in the future (Pierella Karlusich et al., 2022).

Improved global-scale PCC estimates will also include new approaches developed for hyperspectral remote sensing. HPLC is used to develop and validate algorithms that detect pigments and/or PCC from space (Uitz et al., 2015; Chase et al., 2017; Kramer et al., 2022), and the current study presents some encouraging considerations for pigment-based PCC. We find that for many important phytoplankton groups (diatoms, green algae, *Prochlorococcus*, prymnesiophytes), pigments are strongly and positively correlated with PCC from methods with higher taxonomic resolution. This result implies that global PCC assessments should be achievable with the upcoming hyperspectral global ocean color data from NASA’s Plankton Aerosol Cloud and ocean Ecosystem (PACE) mission, planned for launch in 2024 (Werdell et al., 2019). The hyperspectral capability of PACE’s Ocean Color Instrument (OCI) will allow more detailed decomposition of pigment types (Wolanin et al., 2016; Kramer et al., 2022).

If accessory pigments can be accurately modeled from satellite measurements and the comparisons between pigments and amplicon sequencing or IFCB datasets continues on broader spatiotemporal scales, more comprehensive relationships can be developed between pigments and phytoplankton groups determined via other methods throughout the global ocean (Catlett et al., 2022; Chase et al., 2022). By comparing performance across methods, we also encourage consistency in sampling approaches, laboratory analysis, and method development. Including more PCC data types can improve estimates of PCC so long as those data are quality controlled and provide useful information. There are still needs for improvement in many of these comparisons among pigment-based PCC and other methods. For instance, dinoflagellates are an important phytoplankton group (particularly in coastal regions, where they may form toxic blooms), but changes in their relative abundance was not well exclained using Perid/Tchla in the current study. Datasets that have collected samples across a broader range in biomass and under varying physical and biogeochemical conditions, including both coastal (Lin et al., 2019; Catlett et al., 2022) and open ocean systems (Chase et al., 2022) will be ideal for further comparison.

The results shown here are also highly dependent on the relatively small datasets used in our analyses. The relationships among methods will vary based on the region and scale of the comparison. In the datasets used here, certain groups were represented in high relative abundances (e.g., diatoms throughout, *Prochlorococcus* in the NAAMES dataset), resulting in strong relationships among methods despite different sampling approaches. Similar results were found in the West Antarctic Peninsula, where high relative contributions from cryptophytes, diatoms, and prymnesiophytes resulted in significant positive relationships between pigments and 18S (Lin et al., 2019), and in the Neuse River Estuary, where relatively high chlorophyte sequence abundances from 18S correlated well with pigments (Gong et al., 2020). Alternately, in regions with highly dynamic phytoplankton communities and year-round monitoring, the relationships between pigments and other PCC methods were not as clearly defined, due in part to the variability in accessory pigment composition and concentration across the seasonal cycle (Catlett et al., 2022). The capacity of pigments to separate different phytoplankton groups is similarly very dataset-dependent (Kramer and Siegel, 2019), with more and different groups occurring on local scales than global scales. As more high-quality data are collected to further consider the relationships between pigments and other, higher-resolution PCC methods, the scope and scale of those data will necessarily impact the results.

Ultimately, a comprehensive understanding of global surface ocean PCC is essential for better describing relationship between the ocean and global climate, the strength of the biological pump, changes to marine food webs over time, and the cycling of nutrients throughout the oceans. Constraining PCC information from satellites and from discrete water samples is an important and necessary step toward this broader goal. Each PCC method provides one lens through which to view phytoplankton communities, but each view is subject to the constraints of the method. The results shown here suggest that PCC methods provide more information when they are combined and that complementary methods with varied strengths and limitations should be considered wherever possible to provide the most comprehensive understanding of PCC.

## 5. Data availability statement

- HPLC pigments, 18S, 16S, and EXPORTS IFCB data are on SeaBASS: https://seabass.gsfc.nasa.gov/experiment/NAAMES and https://seabass.gsfc.nasa.gov/cruise/EXPORTSNP.
- NAAMES IFCB data are available on EcoTaxa: https://ecotaxa.obs-vlfr.fr.
- Code for IFCB image analysis can be found at: https://github.com/OceanOptics/ifcb-tools (NAAMES) and https://github.com/hsosik/ifcb-analysis (EXPORTS).
- Code for 16S data prep and taxonomic assignment can be found at: https://www.github.com/lbolanos32/NAAMES_2020.
- Code for 18S data prep and taxonomic assignment can be found at: https://github.com/sashajane19/PCCmethods.

## 6. Acknowledgements and funding

- Thank you to Brian VerWey for support with collecting FCM data.
- Thank you to all scientists and technicians on NAAMES and EXPORTS, and to the captains and crews of R/V *Atlantis* and R/V *Sally Ride*.
- Alyson Santoro and Mark Brzezinski provided very helpful edits and comments on an earlier draft of this work.
- Thank you to the Oregon State University Center for Quantitative Life Sciences (16S) and UC Davis Genome Center (18S) for processing the amplicon sequencing data.
- Support for this work came from NASA via the NAAMES and EXPORTS programs (grants NNX15AE72G and 80NSSC17K0692 to DAS; NNX15AAF30G and 80NSSC17K0568 to MJB; NNX15AE67G to EB; 80NSSC17K0654 to CR and HMS); The Simons Foundation (grant 561126 to HMS); Woods Hole Oceanographic Institution’s Ocean Twilight Zone Project, funded as part of The Audacious Project housed at TED (support to HMS, EEP, ETC); the National Defense Science and Engineering Graduate (NDSEG) fellowship through ONR, which supported SJK for 3 years of this work; and NASA PACE grant 80NSSC20M0226 to DAS, which supported SJK for one year of this work.
- We declare no conflicts of interest.

## 7. Supplemental Information

**Figure S1.**
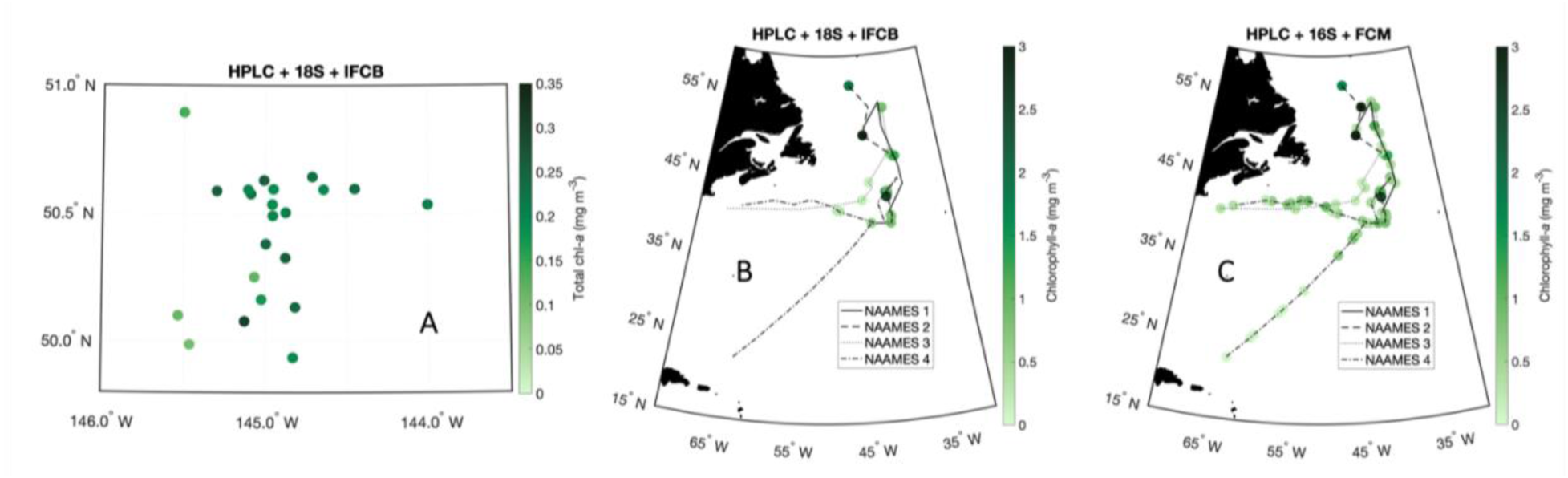
Maps of sampling locations, colored by HPLC total chlorophyll-*a* concentrations, for (A) EXPORTS HPLC + 18S + IFCB, (B) NAAMES HPLC + 18S + IFCB, and (C) NAAMES HPLC + 16S + FCM.

**Figure S2.**
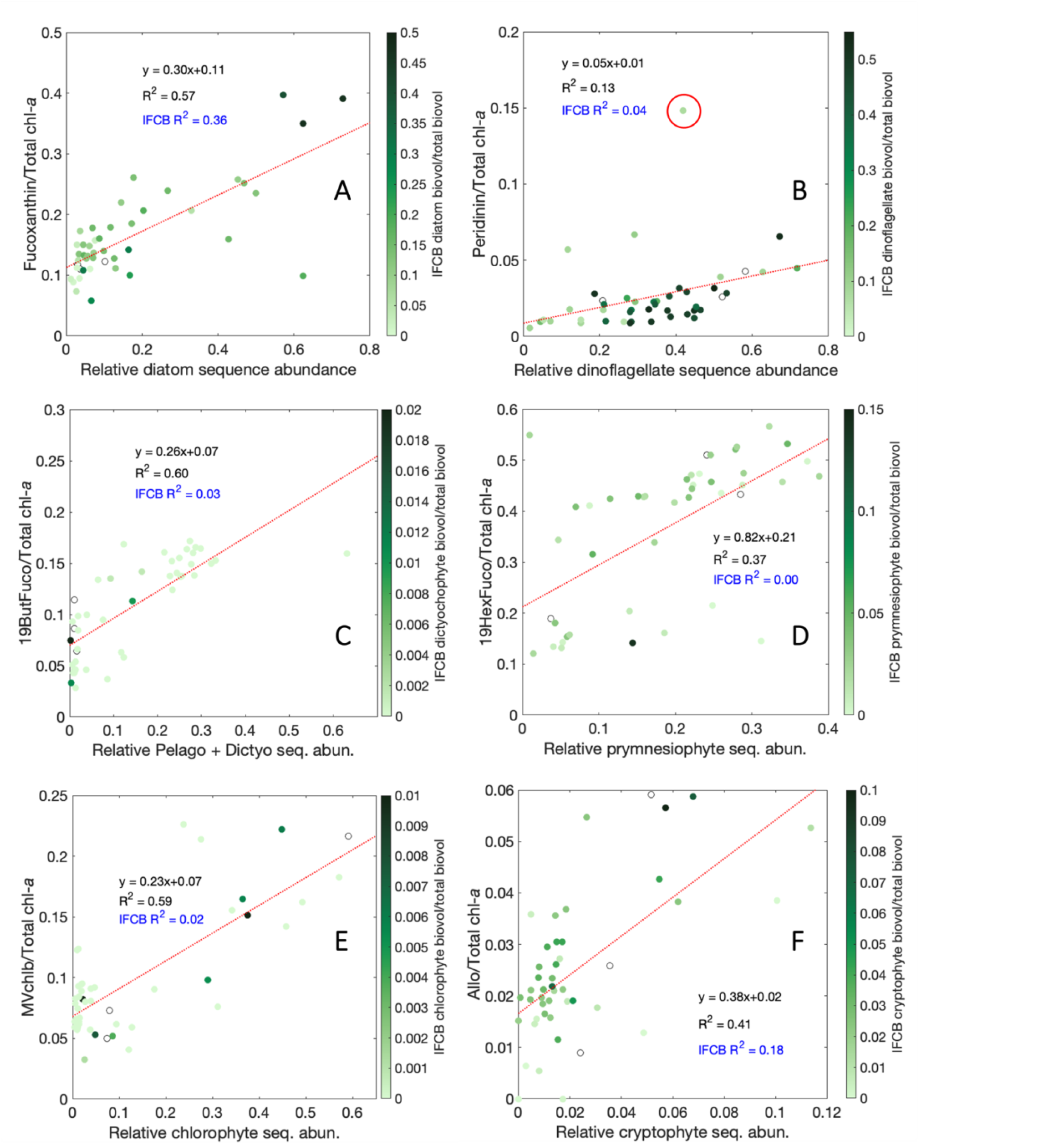
Relationships between relative pigment concentrations (normalized to Tchla) and relative sequence abundances for (A) Fuco and diatoms, (B) Perid and dinoflagellates, (C) 19ButFuco and pelagophytes plus dictyochophytes, (D) 19HexFuco and prymnesiophytes, (E) MVchlb and chlorophytes, and (F) Allo and cryptophytes. All samples are colored by the relative fraction of IFCB biovolume concentration for the corresponding group. Blue text refers to the linear fit for the relative pigment concentration and relative IFCB biovolume concentration (Figure S3). The red circle in (B) denotes a notable outlier.

**Figure S3.**
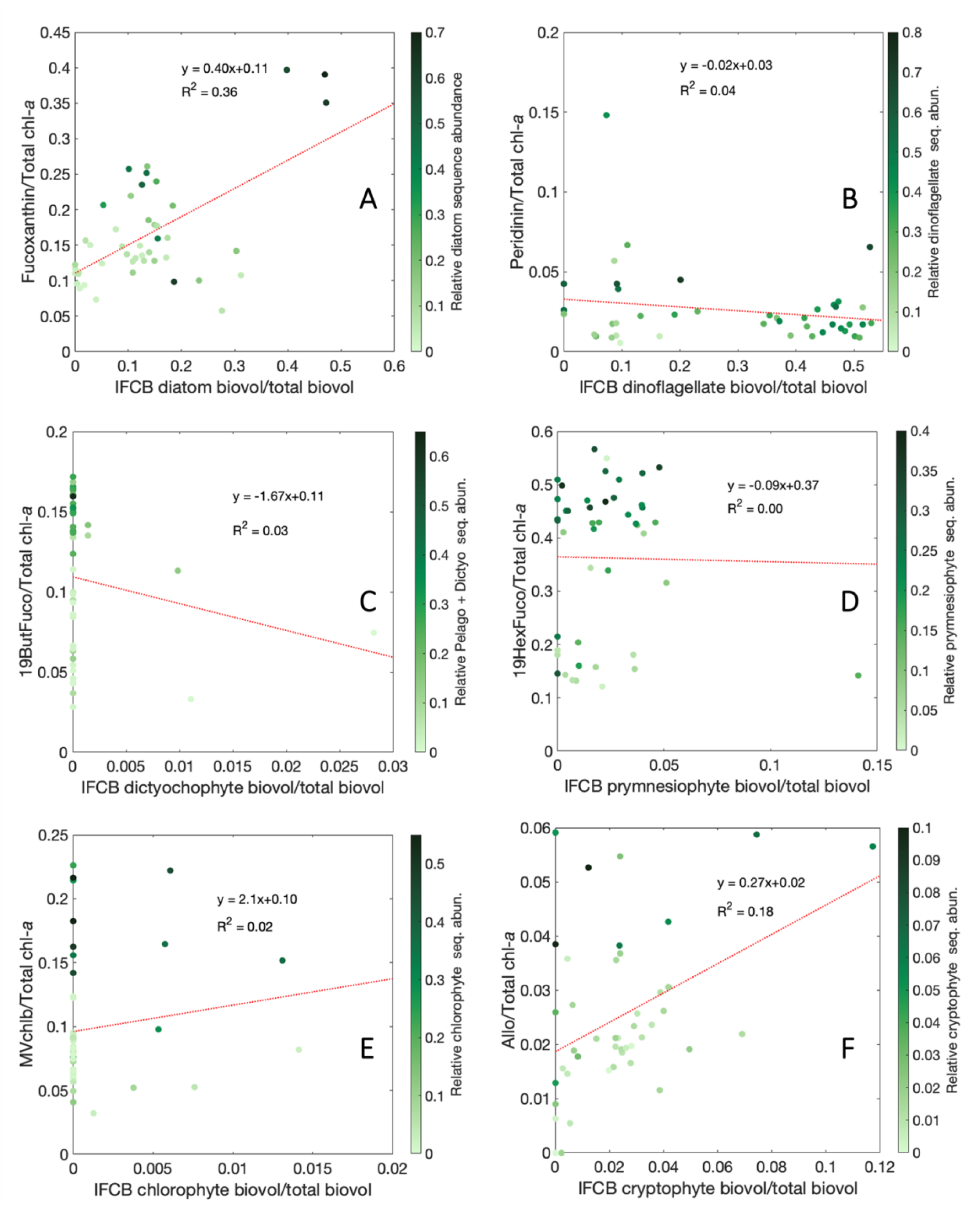
Relationships between relative pigment concentrations (normalized to Tchla) and relative biovolume fractions for (A) Fuco and diatoms, (B) Perid and dinoflagellates, (C) 19ButFuco and dictyochophytes, (D) 19HexFuco and prymnesiophytes, (E) MVchlb and chlorophytes, and (F) Allo and cryptophytes. All samples are colored by the relative fraction of 18S sequence abundances for the corresponding group.

**Figure S4.**
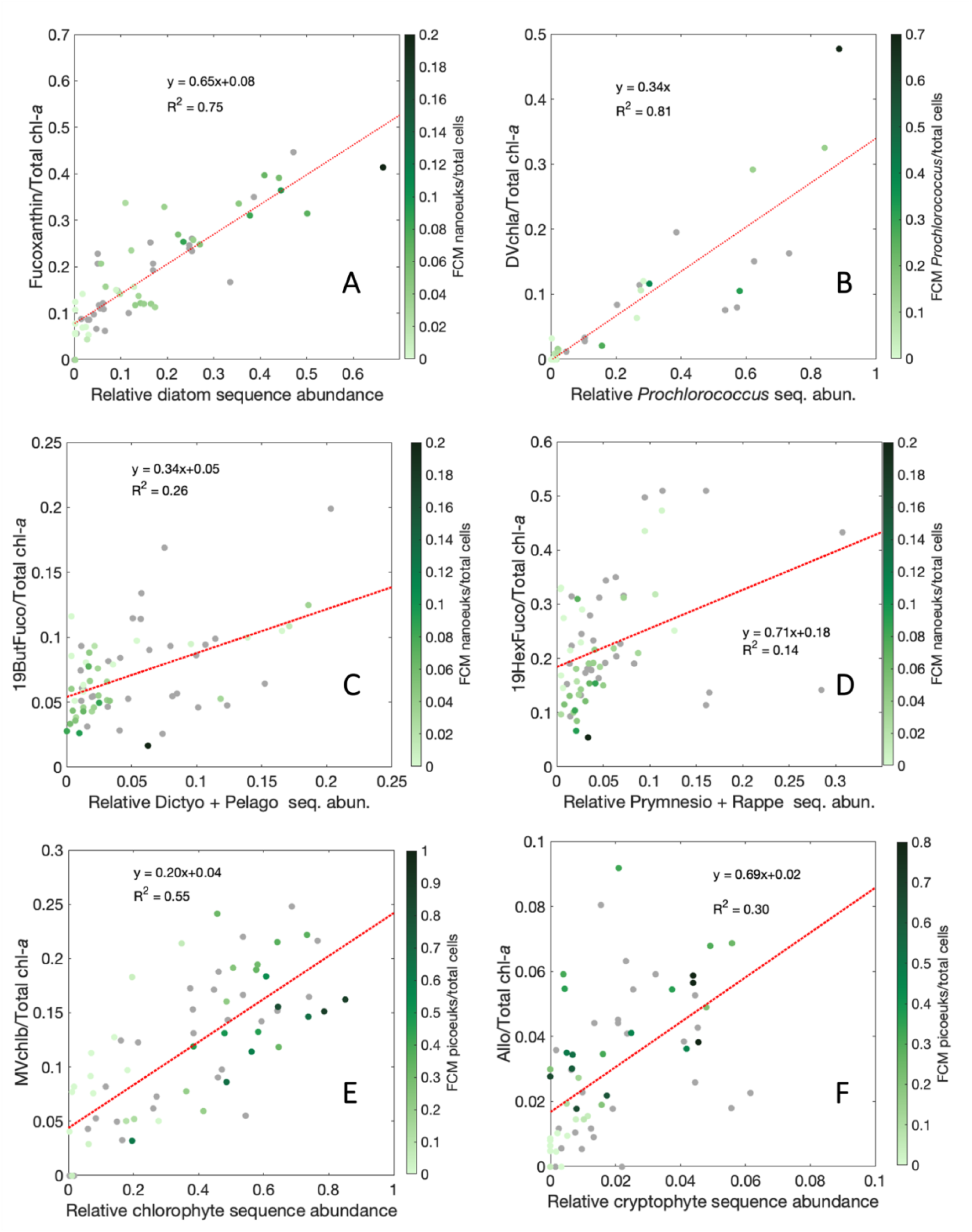
Relationships between relative pigment concentrations and relative sequence abundances for (A) Fuco and diatoms, (B) 19HexFuco and prymnesiophytes plus rappemonads, (C) Allo and cryptophyes, (D) 19ButFuco and dictyochophytes plus pelagophytes, (E) MVchlb and chlorophytes, and (F) DVchla and *Prochlorococcus*. All samples are colored by the relative FCM biovolume concentration that was determined to be most appropriate for that phytoplankton group. Gray dots represent samples for which there was not a FCM matchup.

**Figure S5.**
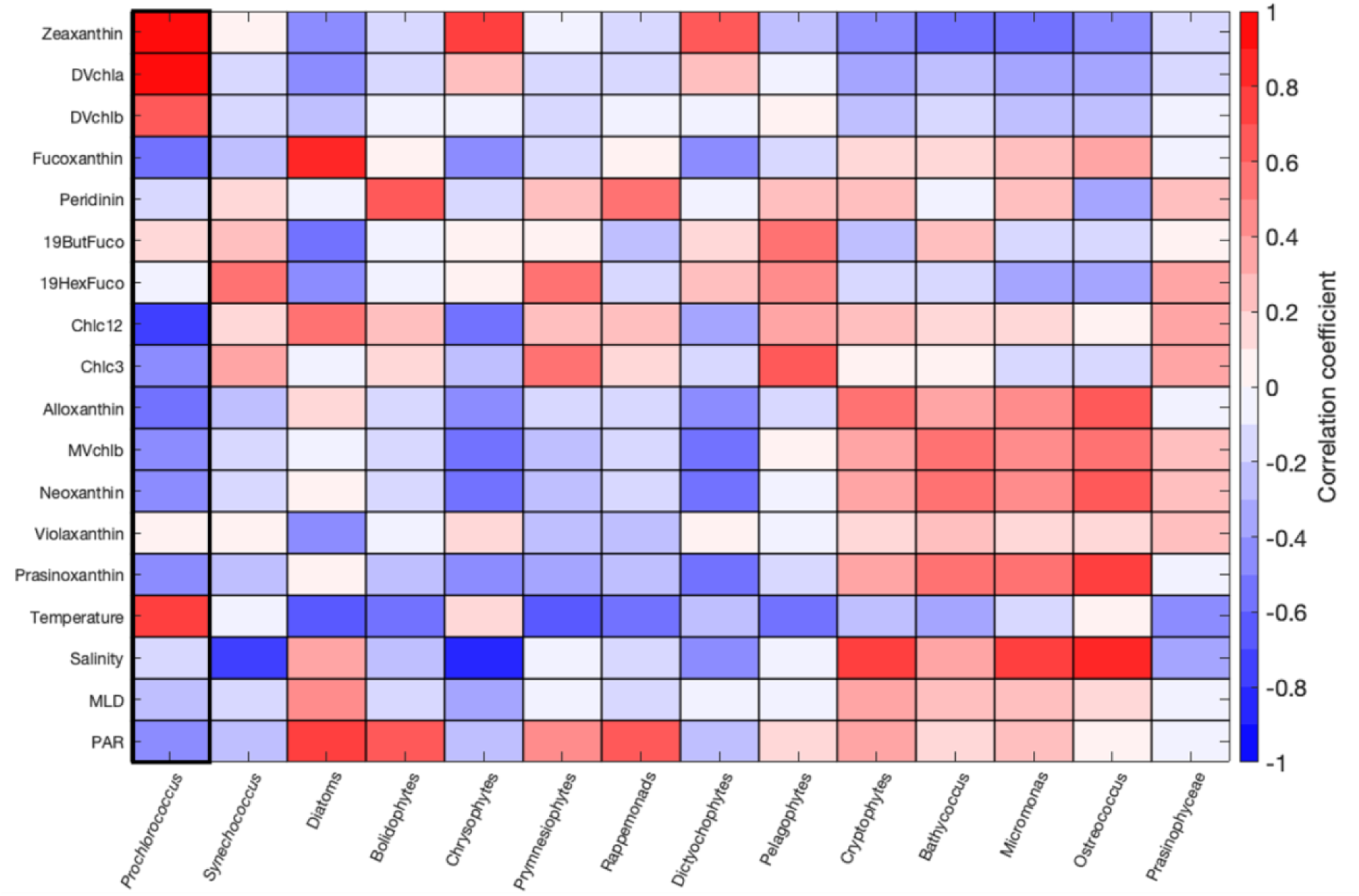
Pearson’s correlation coefficient (R) between relative pigment concentrations and environmental variables (temperature, salinity, MLD, PAR) and relative sequence abundances from 16S.

**Figure S6.**
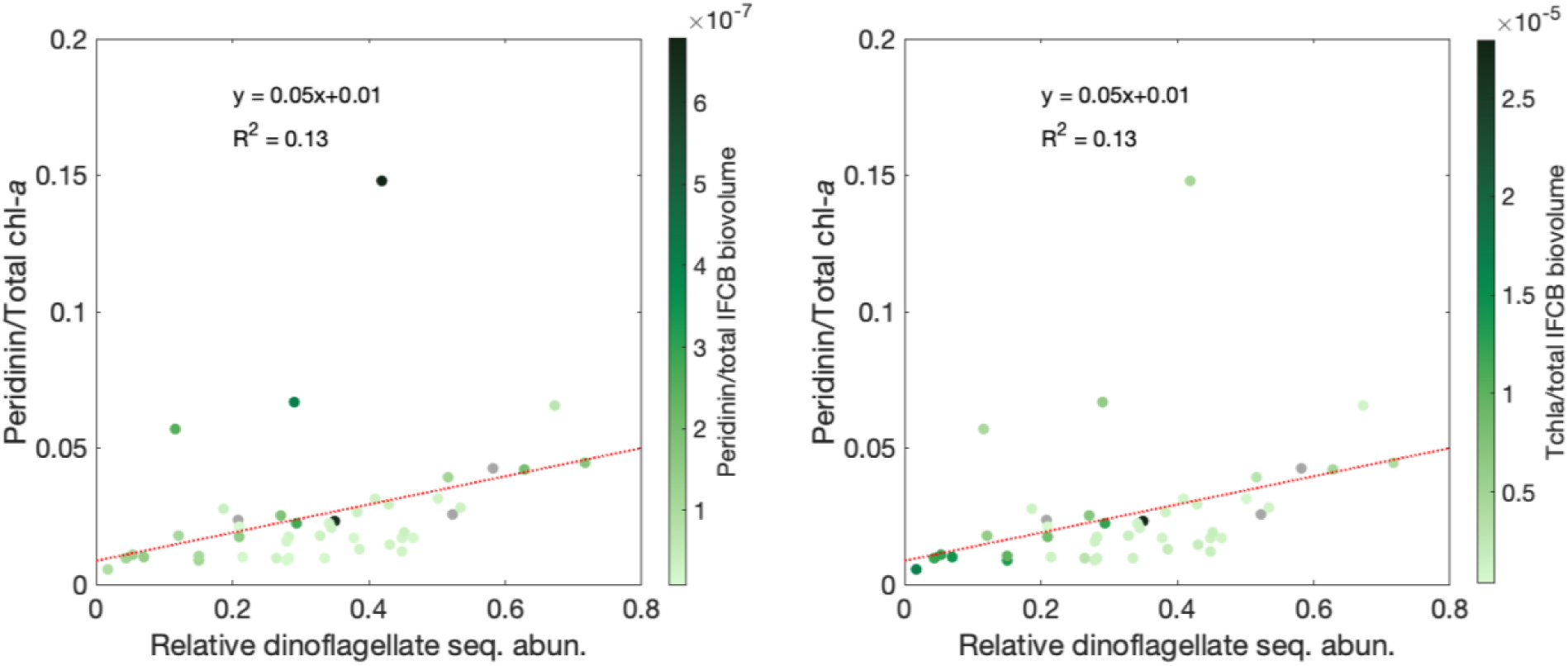
Perid/Tchla vs. relative dinoflagellate sequence abundance colored by (A) Perid concentration per IFCB biovolume concentration and (B) Tchla per IFCB biovolume concentration.

## 8. References

Abad, D., Albaina, A., Aguirre, M., Laza-Martínez, A., Uriarte, I., Iriarte, A., et al. (2016). Is metabarcoding suitable for estuarine plankton monitoring? A comparative study with microscopy. Marine Biology, 163(149), 1–13. 10.1007/s00227-016-2920-0

Adl, S. M., Bass, D., Lane, C. E., Lukeš, J., Schoch, C. L., Smirnov, A., et al. (2019). Revisions to the classification, nomenclature, and diversity of eukaryotes. Journal of Eukaryotic Microbiology, 66(1), 4–119.

Anderson, D. M., Cembella, A. D., & Hallegraeff, G. M. (2012). Progress in Understanding Harmful Algal Blooms: Paradigm Shifts and New Technologies for Research, Monitoring, and Management. Annual Review of Marine Science, 4(1), 143–176. 10.1146/annurev-marine-120308-081121

Behrenfeld, M. J. (2014). Climate-mediated dance of the plankton. Nature Climate Change, 4, 880–887. 10.1038/NCLIMATE2349

Behrenfeld, M. J., Moore, R. H., Hostetler, C. A., Graff, J. R., Gaube, P., Russell, L. M., et al. (2019). The North Atlantic Aerosol and Marine Ecosystem Study (NAAMES): Science motive and mission overview. Frontiers in Marine Science, 6(122), 1–25. 10.3389/fmars.2019.00122

Behrenfeld, M. J., Brooks, S. D., Gaube, P., & Mojica, K. D. A. (n.d.). Unraveling mechanisms underlying annual plankton blooms in the North Atlantic and their implications for biogenic aerosol properties and cloud formation. Frontiers in Marine Science, 8, 6–11. https://www.doi.org/10.3389/fmars.2021.764035

Bolaños, L. M., Karp-Boss, L., Choi, C. J., Worden, A. Z., Graff, J. R., Haëntjens, N., et al. (2020). Small phytoplankton dominate western North Atlantic biomass. The ISME Journal, 14, 1663–1674. 10.1038/S21396-020-0636-0

Bolaños, L. M., Choi, C. J., Worden, A. Z., Baetge, N., Carlson, C. A., & Giovannoni, S. (2021). Seasonality of the Microbial Community Composition in the North Atlantic. Frontiers in Marine Science, 8. 10.3389/fmars.2021.624164

Boss, E., Waite, A., Muller-Karger, F., Yamazaki, H., Wanninkhof, R., Uitz, J., et al. (2018). Beyond Chlorophyll Fluorescence: The Time is Right to Expand Biological Measurements in Ocean Observing Programs. Limnology and Oceanography Bulletin, 27(3), 89–90. 10.1002/lob.10243

Bracher, A., Vountas, M., Dinter, T., Burrows, J. P., Röttgers, R., & Peeken, I. (2009). Quantitative observation of cyanobacteria and diatoms from space using PhytoDOAS on SCIAMACHY data. Biogeosciences, 6, 751–764. https://doi.org/www.biogeosciences.net/6/751/2009/

Bracher, A., Bouman, H. A., Brewin, R. J. W., Bricaud, A., Brotas, V., Ciotti, Á. M., et al. (2017). Obtaining phytoplankton diversity from ocean color: A scientific roadmap for future development. Frontiers in Marine Science, 4, 1–15. 10.3389/fmars.2017.00055

Brosnahan, M. L., Velo-Suárez, L., Ralston, D. K., Fox, S. E., Sehein, T. R., Shalapyonok, A., et al. (2015). Rapid growth and concerted sexual transitions by a bloom of the harmful dinoflagellate Alexandrium fundyense (Dinophyceae). Limnology and Oceanography, 60(6), 2059–2078. 10.1002/lno.10155

Callahan, B. J., McMurdie, P. J., Rosen, M. J., Han, A. W., Johnson, A. J. A., & Holmes, S. P. (2016). DADA2: high-resolution sample inference from Illumina amplicon data. Nature Methods, 13(581), 581–583. 10.1038/nMeth.3869

Campbell, L., Olson, R. J., Sosik, H. M., Abraham, A., Henrichs, D. W., Hyatt, C. J., & Buskey, E. J. (2010). First harmful *Dinophysis* (Dinophyceae, Dinophysiales) bloom in the US is revealed by automated imaging flow cytometry. Journal of Phycology, 46(1), 66–75. 10.1111/j.1529-8817.2009.00791.x

Campbell, L., Gaonkar, C. C., & Henrichs, D. W. (2022). Chapter 5 - Integrating imaging and molecular approaches to assess phytoplankton diversity. In L. A. Clementson, R. S. Eriksen, & A. Willis (Eds.), Advances in Phytoplankton Ecology (pp. 159–190). Elsevier. 10.1016/B978-0-12-822861-6.00013-3

Caron, D. A., Countway, P. D., Jones, A. C., Kim, D. Y., & Schnetzer, A. (2012). Marine Protistan Diversity. Annual Reviews in Marine Science, 4, 467–493. 10.1146/annurev-marine-120709-142802

Caron, D. A., & Hu, S. K. (2019). Are We Overestimating Protistan Diversity in Nature? Trends in Microbiology, 27(3), 197–205. 10.1016/j.tim.2018.10.009

Catlett, D., Matson, P. G., Carlson, C. A., Wilbanks, E. G., Siegel, D. A., & Iglesias-Rodriguez, M. D. (2020). Evaluation of accuracy and precision in an amplicon sequencing workflow for marine protist communities. Limnology and Oceanography: Methods, 18, 20–40. 10.1002/lom3.10343

Catlett, D., Son, K., & Liang, C. (2021a). ensembleTax: an R package for determinations of ensemble taxonomic assignments of phylogenetically-informative marker gene sequences. PeerJ, 9(e11865). 10.7717/peerj.11865

Catlett, D., Siegel, D. A., Simons, R. D., Guillocheau, N., Henderikx-Freitas, F., & Thomas, C. S. (2021b). Diagnosing seasonal to multi-decadal phytoplankton group dynamics in a highly productive coastal ecosystem. Progress in Oceanography, 197, 102637. 10.1016/j.pocean.2021.102637

Catlett, D. S., & Siegel, D. A. (2018). Phytoplankton Pigment Communities Can be Modeled Using Unique Relationships With Spectral Absorption Signatures in a Dynamic Coastal Environment. Journal of Geophysical Research: Oceans, 123, 246–264. 10.1002/2017JC013195

Catlett, D., Siegel, D. A., Matson, P. G., Wear, E. K., Carlson, C. A., Lankiewicz, T. S., & Iglesias-Rodriguez, M. D. (2022). Integrating phytoplankton pigment and DNA meta-barcoding observations to determine phytoplankton composition in the coastal ocean. Limnology and Oceanography, 1–16. 10.1002/lno.12274

Cetinić, I., Poulton, N., & Slade, W. H. (2016). Characterizing the phytoplankton soup: pump and plumbing effects on the particle assemblage in underway optical seawater systems. Optics Express, 24(18), 20703–20715. 10.1364/OE.24.020703

Chase, A. P., Boss, E., Zaneveld, R., Bricaud, A., Claustre, H., Ras, J., et al. (2013). Decomposition of in situ particulate absorption spectra. Methods in Oceanography, 7, 110–124. 10.1016/j.mio.2014.02.002

Chase, A. P., Boss, E., Cetinić, I., & Slade, W. (2017). Estimation of Phytoplankton Accessory Pigments from Hyperspectral Reflectance Spectra: Toward a Global Algorithm. Journal of Geophysical Research: Oceans, 122, 1–19. 10.1002/2017JC012859

Chase, A. P., Boss, E. S., Haëntjens, N., Culhane, E., Roesler, C., & Karp-Boss, L. (2022). Plankton Imagery Data Inform Satellite-Based Estimates of Diatom Carbon. Geophysical Research Letters, 49(13), e2022GL098076. 10.1029/2022GL098076

Chase, A. P., Kramer, S. J., Haëntjens, N., Boss, E. S., Karp-Boss, L., Edmondson, M., & Graff, J. R. (2020). Evaluation of diagnostic pigments to estimate phytoplankton size classes. Limnology and Oceanography: Methods, 18(10), 570–584. https://www.doi.org/10.1002/lom3.10385

Coupel, P., Matsuoka, A., Ruiz-Pino, D., Gosselin, M., Marie, D., Tremblay, J.-É., & Babin, M. (2015). Pigment signatures of phytoplankton communities in the Beaufort Sea. Biogeosciences, 12, 991–1006. 10.5194/bg-12-991-2015

Della Penna, A., & Gaube, P. (2019). Overview of (sub)mesoscale ocean dynamics for the NAAMES field program. Frontiers in Marine Science, 6(384), 1–7. 10.3389/fmars.2019.00384

Durkin, C., Cetinić, I., Estapa, M. L., Ljubešić, Z., Mucko, M., Neeley, A., & Omand, M. M. (2022). Tracing the path of carbon export in the ocean though DNA sequencing of individual sinking particles. The ISME Journal, 1–11. 10.1038/S21396-022-01239-2

Estapa, M. L., Siegel, D. A., Buesseler, K. O., Stanley, R. H. R., Lomas, M. W., & Nelson, N. B. (2015). Decoupling of net community and export production on submesoscales in the Sargasso Sea. Global Biogeochemical Cycles, 29(8), 1266–1282. 10.1002/2014GB004913

Gloor, G. B., Macklaim, J. M., Pawlowsky-Glahn, V., & Egozcue, J. J. (2017). Microbiome Datasets Are Compositional: And This Is Not Optional. Frontiers in Microbiology, 8. 10.3389/fmicb.2017.02224

Godhe, A., Asplund, M. E., Härnström, K., Saravanan, V., Tyagi, A., & Karunasagar, I. (2008). Quantification of diatom and dinoflagellate biomasses in coastal marine seawater samples by real-time PCR. Applied and Environmental Microbiology, 74(23), 7174–7182. 10.1128/AEM.01298-08

Gong, W., Hall, N., Paerl, H., & Marchetti, A. (2020). Phytoplankton composition in a eutrophic estuary: Comparison of multiple taxonomic approaches and influence of environmental factors. Environmental Microbiology, 22(11), 4718–4731. 10.1111/1462-2920.15221

Graff, J. R., & Behrenfeld, M. J. (2018). Photoacclimation responses in subarctic Atlantic phytoplankton following a natural mixing-restratification event. Frontiers in Marine Science, 5, 1–11. 10.3389/fmars.2018.00209

Gu, Z., Gu, L., Eils, R., Schlesner, M., & Brors, B. (2014). circlize implements and enhances circular visualization in R. Bioinformatics, 30(19), 2811–2812. 10.1093/bioinformatics/btu393

Guidi, L., Legendre, L., Reygondeau, G., Uitz, J., Stemmann, L., & Henson, S. A. (2015). A new look at ocean carbon remineralization for estimating deepwater sequestration. Global Biogeochemical Cycles, 29(7), 1044–1059. 10.1002/2014GB005063

Guidi, L., Chaffron, S., Bittner, L., Eveillard, D., Larhlimi, A., Roux, S., et al. (2016). Plankton networks driving carbon export in the oligotrophic ocean. Nature, 532, 465–470. 10.1038/nature16942

Guillou, L., Bachar, D., Audic, S., Bass, D., Berney, C., Bittner, L., et al. (2013). The Protist Ribosomal Reference database (PR2): a catalog of unicellular eukaryote Small Sub-Unit rRNA sequences with curated taxonomy. Nucleic Acids Research, 41(D1), D597–D604. 10.1093/nar/gks1160

Haëntjens, N., Boss, E. S., Graff, J. R., Chase, A. P., & Karp-Boss, L. (2022). Phytoplankton size distributions in the western North Atlantic and their seasonal variability. Limnology and Oceanography, 67(8), 1865–1878. 10.1002/lno.12172

Henriksen, P., Riemann, B., Kaas, H., Sørenson, H. M., & Sørenson, H. L. (2002). Effects of nutrient-limitation and irradiance on marine phytoplankton pigments. Journal of Plankton Research, 24(9), 835–858. 10.1093/plankt/24.9.835

Hooker, S. B., Clementson, L., Thomas, C. S., Schlüter, L., Allerup, M., Ras, J., et al. (2012). The Fifth SeaWiFS HPLC Analysis Round-Robin Experiment (SeaHARRE-5) (NASA Technical Reports) (pp. 1–108). Greenbelt, Maryland: NASA Goddard Space Flight Center.

Hunter-Cevera, K. R., Neubert, M. G., Olson, R. J., Solow, A. R., Shalapyonok, A., & Sosik, H. M. (2016). Physiological and ecological drivers of early spring blooms of a coastal phytoplankter. Science, 354(6310), 326–329. 10.1126/science.aaf8536

Hunter-Cevera, K. R., Hamilton, B. R., Neubert, M. G., & Sosik, H. M. (2021). Seasonal environmental variability drives microdiversity within a coastal Synechococcus population. Environmental Microbiology, 23(8), 4689–4705. 10.1111/1462-2920.15666

Irigoien, X., Meyer, B., Harris, R. P., & Harbour, D. S. (2004). Using HPLC pigment analysis to investigate phytoplankton taxonomy: the importance of knowing your species. Helgoland Marine Research, 58, 77–82. 10.1007/s10152-004-0171-9

Jeffrey, S. W., Wright, S. W., & Zapata, M. (2011). Microalgal classes and their signature pigments. In S. Roy, C. A. Llewellyn, E. S. Egeland, & G. Johnsen (Eds.), Phytoplankton Pigments: Characterization, Chemotaxonomy, and Application in Oceanography (pp. 3– 77). Cambridge, United Kingdom: Cambridge University Press.

Johnson, Z. I., & Martiny, A. C. (2015). Techniques for quantifying phytoplankton biodiversity. The Annual Review of Marine Science, 7, 299–324. 10.1146/annurev-marine-010814-015902

Karlson, B., Godhe, A., Cusack, C., & Bresnan, E. (2010). Introduction to methods for quantitative phytoplankton analysis. Microscopic and Molecular Methods for Quantitative Phytoplankton Analysis, 5. Paris, UNESCO.

Kawachi, M., Nakayama, T., Kayama, M., Nomura, M., Miyashita, H., Bojo, O., et al. (2021). Rappemonads are haptophyte phytoplankton. Current Biology, 31(11), 2395–2403.e4. 10.1016/j.cub.2021.03.012

Kramer, S. J., & Siegel, D. A. (2019). How can phytoplankton pigments be best used to characterize surface ocean phytoplankton groups for ocean color remote sensing algorithms? Journal of Geophysical Research: Oceans, 124, 7557–7574. 10.1029/2019JC015604

Kramer, S. J., Siegel, D. A., & Graff, J. R. (2020a). Phytoplankton community composition determined from co-variability among phytoplankton pigments from the NAAMES field campaign. Frontiers in Marine Science, 7, 1–15. 10.3389/fmars.2020.00215

Kramer, S. J., Bisson, K. M., & Fischer, A. D. (2020b). Observations of phytoplankton community composition in the Santa Barbara Channel during the Thomas Fire. Journal of Geophysical Research: Oceans, 125(12), 1–16. 10.1029/2020JC016851

Kramer, S. J., Siegel, D. A., Maritorena, S., & Catlett, D. (2022). Modeling surface ocean phytoplankton pigments from hyperspectral remote sensing reflectance on global scales. Remote Sensing of Environment, 270, 112879. 10.1016/j.rse.2021.112879

Kuwata, A., Yamada, K., Ichinomiya, M., Yoshikawa, S., Tragin, M., Vaulot, D., & Lopes dos Santos, A. (2018). Bolidophyceae, a Sister Picoplanktonic Group of Diatoms – A Review. Frontiers in Marine Science, 5. 10.3389/fmars.2018.00370

Laney, S. R., & Sosik, H. M. (2014). Phytoplankton assemblage structure in and around a massive under-ice bloom in the Chukchi Sea. The Phytoplankton Megabloom beneath Arctic Sea Ice: Results from the ICESCAPE Program, 105, 30–41. 10.1016/j.dsr2.2014.03.012

Legendre, L. (1990). The significance of microalgal blooms for fisheries and for the export of particulate organic carbon in oceans. Journal of Plankton Research, 12(4), 681–699. 10.1093/plankt/12.4.681

Li, Q., Edwards, K. F., Schvarcz, C. R., & Steward, G. F. (2022). Broad phylogenetic and functional diversity among mixotrophic consumers of Prochlorococcus. The ISME Journal. 10.1038/S21396-022-01204-z

Lima-Mendez, G., Faust, K., Henry, N., Decelle, J., Colin, S., Carcillo, F., et al. (2015). Determinants of community structure in the global plankton interactome. Science, 348(6237), 1262073. 10.1126/science.1262073

Lin, S. (2011). Genomic understanding of dinoflagellates. The Genome Organisation of Eukaryotic Microbes, 162(6), 551–569. 10.1016/j.resmic.2011.04.006

Lin, Y., Gifford, S., Ducklow, H., Schofield, O., & Cassar, N. (2019). Towards quantitative microbiome community profiling using internal standards. Applied and Environmental Microbiology, 85(5), 1–14. 10.1128/AEM.02634-18

Lombard, F., Boss, E., Waite, A. M., Vogt, M., Uitz, J., Stemmann, L., et al. (2019). Globally consistent quantitative observations of planktonic ecosystems. Frontiers in Marine Science, 6, 1–21. 10.3389/fmars.2019.00196

Mackey, M. D., Mackey, D. J., Higgins, H. W., & Wright, S. W. (1996). CHEMTAX - a program for estimating class abundances from chemical markers: application to HPLC measurements of phytoplankton. Marine Ecology Progress Series, 144, 265–283. 10.3354/meps144265

Martiny, A. C., Pham, C. T. A., Primeau, F. W., Vrugt, J. A., Moore, J. K., Levin, S. A., & Lomas, M. W. (2013). Strong latitudinal patterns in the elemental ratios of marine plankton and organic matter. Nature Geoscience, 6(4), 279–283. 10.1038/ngeo1757

Menden-Deuer, S., & Lessard, E. J. (2000). Carbon to volume relationships for dinoflagellates, diatoms, and other protist plankton. Limnology and Oceanography, 45(3), 569–579. 10.4319/lo.2000.45.3.0569

Moberg, E. A., & Sosik, H. M. (2012). Distance maps to estimate cell volume from two-dimensional plankton images. Limnology and Oceanography: Methods, 10, 278–288. 10.4319/lom.2012.10.278

Murali, A., Bhargava, A., & Wright, E. S. (2018). IDTAXA: a novel approach for accurate taxonomic classification of microbiome sequences. Microbiome, 6(140). 10.1186/S20168-018-0521-5

Nardelli, S. C., Gray, P. C., Stammerjohn, S. E., & Schofield, O. (2023). Characterizing coastal phytoplankton seasonal succession patterns on the West Antarctic Peninsula. Limnology and Oceanography, 1-17. 10.1002/lno.12314

Nascimento, S. M., Purdie, D. A., & Morris, S. (2005). Morphology, toxin composition and pigment content of Prorocentrum lima strains isolated from a coastal lagoon in southern UK. Toxicon, 45(5), 633–649. 10.1016/j.toxicon.2004.12.023

National Academies of Sciences, Engineering, and Medicine. (2022). Nutrient Fertilization. In A Research Strategy for Ocean-based Carbon Dioxide Removal and Sequestration. Washington, DC: The National Academies Press. 10.17226/26278

Nayar, S., & Chou, L. M. (2003). Relative efficiencies of different filters in retaining phytoplankton for pigment and productivity studies. Estuarine, Coastal and Shelf Science, 58(2), 241–248. 10.1016/S0272-7714(03)00075-1

Needham, D. M., & Fuhrman, J. A. (2016). Pronounced daily succession of phytoplankton, archaea and bacteria following a spring bloom. Nature Microbiology, 1–7. 10.1038/NMICROBIOL.2016.5

Neeley, A. R., Lomas, M. W., Mannino, A., Thomas, C., & Vandermeulen, R. (2022). Impact of Growth Phase, Pigment Adaptation, and Climate Change Conditions on the Cellular Pigment and Carbon Content of Fifty-One Phytoplankton Isolates. Journal of Phycology. 10.1111/jpy.13279

Not, F., Latasa, M., Scharek, R., Viprey, M., Karleskind, P., Balagué, V., et al. (2008). Protistan assemblages across the Indian Ocean, with a specific emphasis on the picoeukaryotes. Deep Sea Research Part I: Oceanographic Research Papers, 55(11), 1456–1473. 10.1016/j.dsr.2008.06.007

Olson, R. J., & Sosik, H. M. (2007). A submersible imaging-in-flow instrument to analyze nano- and microplankton: Imaging FlowCytobot. Limnology and Oceanography: Methods, 5(6), 195–203. 10.4319/lom.2007.5.195

Peacock, E. E., Olson, R. J., & Sosik, H. M. (2014). Parasitic infection of the diatom *Guinardia delicatula*, a recurrent and ecologically important phenomenon on the New England Shelf. Marine Ecology Progress Series, 503, 1–10. 10.3354/meps10784

Picheral, M., Colin, S., & Irisson, J.-O. (2017). EcoTaxa, a tool for the taxonomic classification of images. Retrieved from https://ecotaxa.obs-vlfr.fr.

Pierella Karlusich, J. J., Pelletier, E., Zinger, L., Lombard, F., Zingone, A., Colin, S., et al. (2022). A robust approach to estimate relative phytoplankton cell abundances from metagenomes. Molecular Ecology Resources. 10.1111/1755-0998.13592

Quast, C., Pruesse, E., Yilmaz, P., Gerken, J., Schweer, T., Yarza, P., et al. (2012). The SILVA ribosomal RNA gene database project: improved data processing and web-based tools. Nucleic Acid Research, 41(D1), D590–D596. 10.1093/nar/gks1219

Rubinov, M., & Sporns, O. (2010). Complex network measures of brain connectivity: Uses and interpretations. NeuroImage, 52(3). 10.1016/j.neuroimage.2009.10.003

Russakovsky, O., Deng, J., Su, H., Krause, J., Satheesh, S., Ma, S., Huang, Z., Karpathy, A., Khosla, A., Bernstein, M., Berg, A.C., & Fei-Fei, L. (2015). ImageNet large scale visual recognition challenge. International Journal of Computer Vision, 115, 211–252. 10.1007/s11263-015-0816-y

Schlüter, L., Møhlenberg, F., Havskum, H., & Larsen, S. (2000). The use of phytoplankton pigments for identifying and quantifying phytoplankton groups in coastal areas: testing the influence of light and nutrients on pigment/chlorophyll a ratios. Marine Ecology Progress Series, 192, 49–63. 10.3354/meps192049

Siegel, D. A., Westberry, T. K., O’Brien, M. C., Nelson, N. B., Michaels, A. F., Morrison, J. R., et al. (2001). Bio-optical modeling of primary production on regional scales: the Bermuda BioOptics project. Deep Sea Research Part II: Topical Studies in Oceanography, 48(8), 1865–1896. 10.1016/S0967-0645(00)00167-3

Siegel, D. A., Buesseler, K. O., Behrenfeld, M. J., Benitez-Nelson, C. R., Boss, E., Brzezinski, M. A., et al. (2016). Prediction of the Export and Fate of Global Ocean Net Primary Production: The EXPORTS Science Plan. Frontiers in Marine Science, 3. https://www.doi.org/10.3389/fmars.2016.00022

Siegel, D. A., Cetinić, I., Graff, J. R., Lee, C. M., Nelson, N., Perry, M. J., et al. (2021). An operational overview of the EXport Processes in the Ocean from RemoTe Sensing (EXPORTS) Northeast Pacific field deployment. Elementa: Science of the Anthropocene, 9(1). 10.1525/elementa.2020.00107

Siegel, D. A., DeVries, T., Cetinić, I., & Bisson, K. M. (2023). Quantifying the Ocean’s Biological Pump and Its Carbon Cycle Impacts on Global Scales. Annual Review of Marine Science, 15(1), 329–356. 10.1146/annurev-marine-040722-115226

Sommeria-Klein, G., Watteaux, R., Iudicone, D., Bowler, C., & Morlon, H. (2021). Global drivers of eukaryotic plankton biogeography in the sunlit ocean. Science, 374(6567), 594–599. https://www.doi.org/10.1126/science.abb3717

Sosik, H. M., & Olson, R. J. (2007). Automated taxonomic classification of phytoplankton sampled with imaging-in-flow cytometry. Limnology and Oceanography: Methods, 5, 204–216. 10.4319/lom.2007.5.204

Sosik, H. M., Olson, R. J., & Armbrust, E. V. (2010). Flow Cytometry in Phytoplankton Research. In D. J. Suggett, O. Prasil, & M. A. Borowitzka (Eds.), Chlorophyll-a fluorescence in aquatic science: methods and applications. Developments in Applied Phycology 4 (pp. 171–185). Springer.

Sosik, H. M., Sathyendranath, S., Uitz, J., Bouman, H., & Nair, A. (2014). *In situ* Methods of Measuring Phytoplankton Functional Types. In Phytoplankton Functional Types from Space (Vol. 15, pp. 21–38). Dartmouth, Canada.

Stoecker, D. K., Hansen, P. J., Caron, D. A., & Mitra, A. (2017). Mixotrophy in the Marine Plankton. Annual Review of Marine Science, 9(1), 311–335. 10.1146/annurev-marine-010816-060617

Szegedy, C., Wei, L., Jia, Y., Sermanet, P. Reed, S., Anguelov, D., Erhan, D., Vanhoucke, V., & Rabinovich, A. (2015). Going deeper with convolutions. Proceedings of the IEEE Conference on Computer Vision and Pattern Recognition (CVPR), pp. 1–9. 10.1109/CVPR.2015.7298594.

Thibodeau, P. S., Roesler, C. S., Drapeau, S. L., Prabhu Matondkar, S. G., Goes, J. I., & Werdell, P. J. (2014). Locating Noctiluca miliaris in the Arabian Sea: An optical proxy approach. Limnology and Oceanography, 59(6), 2042–2056. 10.4319/lo.2014.59.6.2042

Thompson, P. A., Pesant, S., & Waite, A. M. (2007). Contrasting the vertical differences in the phytoplankton biology of a dipole pair of eddies in the south-eastern Indian Ocean. The Leeuwin Current and Its Eddies, 54(8), 1003–1028. 10.1016/j.dsr2.2006.12.009

Trudnowska, E., Lacour, L., Ardyna, M., Rogge, A., Irisson, J.-O., Waite, A. M., et al. (2021). Marine snow morphology illuminates the evolution of phytoplankton blooms and determines their subsequent vertical export. Nature Communications, 12(2816), 1–13. 10.1038/S21467-021-22994-4

Uitz, J., Stramski, D., Reynolds, R. A., & Dubranna, J. (2015). Assessing phytoplankton community composition from hyperspectral measurements of phytoplankton absortion coefficient and remote-sensing reflectance in open-ocean environments. Remote Sensing of the Environment, 171, 58–74. 10.1016/j.rse.2015.09.027

Vallina, S. M., Cermeno, P., Dutkiewicz, S., Loreau, M., & Montoya, J. M. (2017). Phytoplankton functional diversity increases ecosystem productivity and stability. Ecological Modelling, 361, 184–196. 10.1016/j.ecolmodel.2017.06.020

Van Heukelem, L., & Hooker, S. B. (2011). The importance of a quality assurance plan for method validation and minimizing uncertainties in the HPLC analysis of phytoplankton pigments. In S. Roy, C. A. Llewellyn, E. S. Egeland, & G. Johnsen (Eds.), Phytoplankton Pigments: Characterization, Chemotaxonomy, and Applications in Oceanography (pp. 195–242). Cambridge, United Kingdom: Cambridge University Press.

Van Heukelem, L., & Thomas, C. S. (2001). Computer-assisted high-performance liquid chromatography method development with applications to the isolation and analysis of phytoplankton pigments. Journal of Chromatography A, 910, 10.1016/S0378-4347(00)00603-4.

de Vargas, C., Audic, S., Henry, N., Decelle, J., Mahé, F., Logares, R., et al. (2015). Eukaryotic plankton diversity in the sunlit ocean. Science, 348(6237), 1–11. 10.1126/science.1261605

Vergin, K. L., Beszteri, B., Monier, A., Cameron Thrash, J., Temperton, B., Treusch, A. H., et al. (2013). High-resolution SAR11 ecotype dynamics at the Bermuda Atlantic Time-series Study site by phylogenetic placement of pyrosequences. The ISME Journal, 7(7), 1322– 1332. 10.1038/ismej.2013.32

Werdell, P. J., Behrenfeld, M. J., Bontempi, P. S., Boss, E., Cairns, B., Davis, G. T., et al. (2019). The Plankton, Aerosol, Cloud, ocean Ecosystem (PACE) mission: Status, science, advances. Bulletin of the American Meteorological Society, 1–59. 10.1175/BAMS-D-18-0056.1

Wolanin, A., Soppa, M. A., & Bracher, A. (2016). Investigation of spectral band requirements for improving retrievals of Phytoplankton Functional Types. Remote Sensing, 8(871), 1–21. 10.3390/rs6100871

Yeh, Y.-C., & Fuhrman, J. A. (2022). Contrasting diversity patterns of prokaryotes and protists over time and depth at the San-Pedro Ocean Time series. ISME Communications, 2(1), 36. 10.1038/s43705-022-00121-8

Yilmaz, P., Parfrey, L. W., Yarza, P., Gerken, J., Pruesse, E., Quast, C., et al. (2014). The SILVA and “All-species Living Tree Project (LTP)” taxonomic frameworks. Nucleic Acid Research, 42, D643–D648. 10.1093/nar/gkt1209

Zapata, M, Fraga, S., Rodríguez, F., & Garrido, J. L. (2012). Pigment-based chloroplast types in dinoflagellates. Marine Ecology Progress Series, 465, 33–52. 10.3354/meps09879

Zapata, Manuel, Jeffrey, S. W., Wright, S. W., Rodríguez, F., Garrido, J. L., & Clementson, L. (2004). Photosynthetic pigments in 37 species (65 strains) of Haptophyta: implications for oceanography and chemotaxonomy. Marine Ecology Progress Series, 270, 83–102. 10.3354/meps270083

Zhang, B., & Horvath, S. (2005). A General Framework for Weighted Gene Co-Expression Network Analysis. Statistical Applications in Genetics and Molecular Biology, 4(1), 1–45. 10.2202/1544-6115.1128

Zhang, X., Huot, Y., Bricaud, A., & Sosik, H. M. (2015). Inversion of spectral absorption coefficients to infer phytoplankton size classes, chlorophyll concentration, and detrital matter. Applied Optics, 54(18), 5805–5816. 10.1364/AO.54.005805

Zhou, J., & Ning, D. (2017). Stochastic Community Assembly: Does It Matter in Microbial Ecology? Microbiology and Molecular Biology Reviews : MMBR, 81(4), e00002–17. 10.1128/MMBR.00002-17

Zhu, F., Massana, R., Not, F., Marie, D., & Vaulot, D. (2005). Mapping of picoeucaryotes in marine ecosystems with quantitative PCR of the 18S rRNA gene. FEMS Microbiology Ecology, 52(1), 79–92. 10.1016/j.femsec.2004.10.006

Zubkov, M. V., Sleigh, M. A., Tarran, G. A., Burkill, P. H., & Leakey, R. J. G. (1998). Picoplanktonic community structure on an Atlantic transect from 50°N to 50°S. Deep Sea Research Part I: Oceanographic Research Papers, 45(8), 1339–1355. 10.1016/S0967-0637(98)00015-6

