## Supplemental figures for "Toward a synthesis of phytoplankton community composition methods for global-scale application"

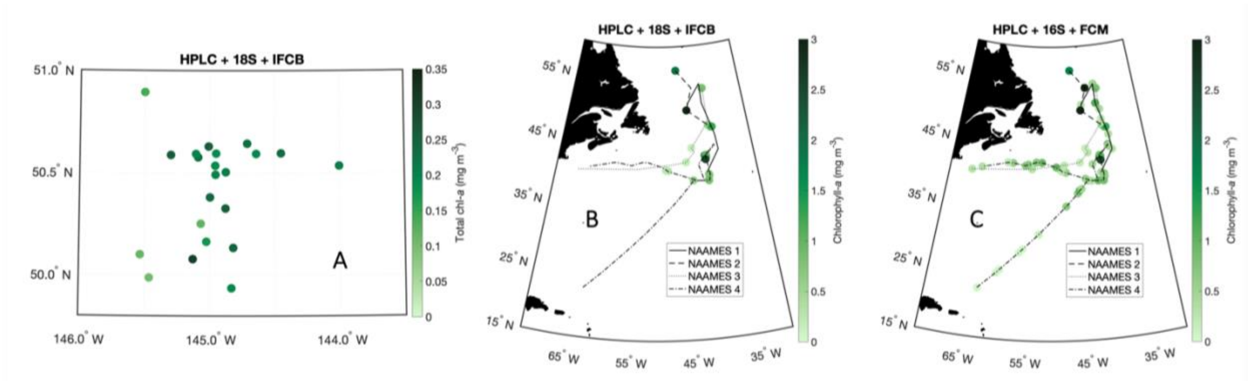


**Figure S1.** Maps of sampling locations, colored by HPLC total chlorophyll-*a* concentrations, for (A) EXPORTS HPLC + 18S + IFCB, (B) NAAMES HPLC + 18S + IFCB, and (C) NAAMES HPLC + 16S + FCM.

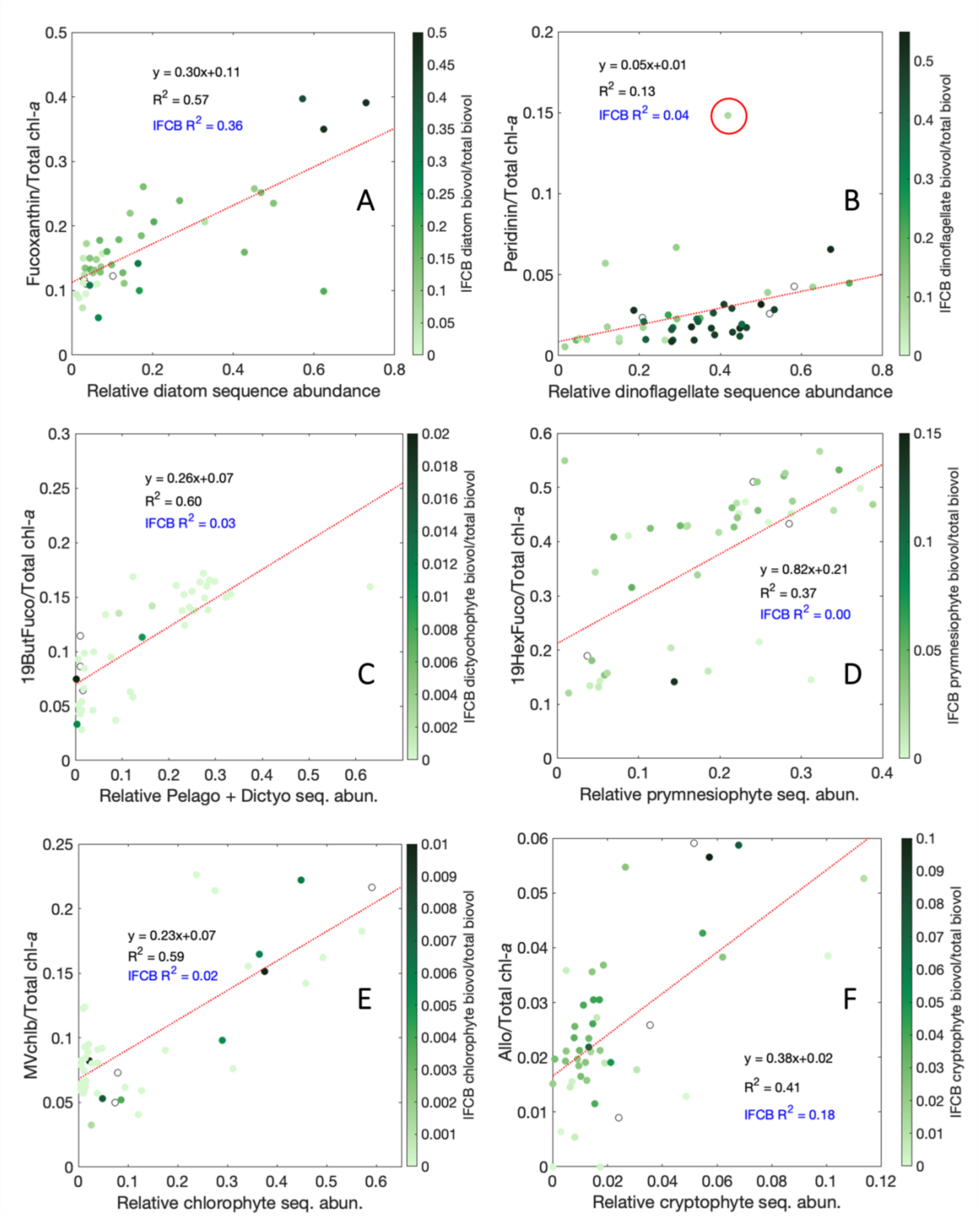


**Figure S2.** Relationships between relative pigment concentrations (normalized to Tchla) and relative sequence abundances for (A) Fuco and diatoms, (B) Perid and dinoflagellates, (C) 19ButFuco and pelagophytes plus dictyochophytes, (D) 19HexFuco and prymnesiophytes, (E) MVchlb and chlorophytes, and (F) Allo and cryptophytes. All samples are colored by the relative fraction of IFCB biovolume concentration for the corresponding group. Blue text refers to the linear fit for the relative pigment concentration and relative IFCB biovolume concentration (Figure S3). The red circle in (B) denotes a notable outlier.

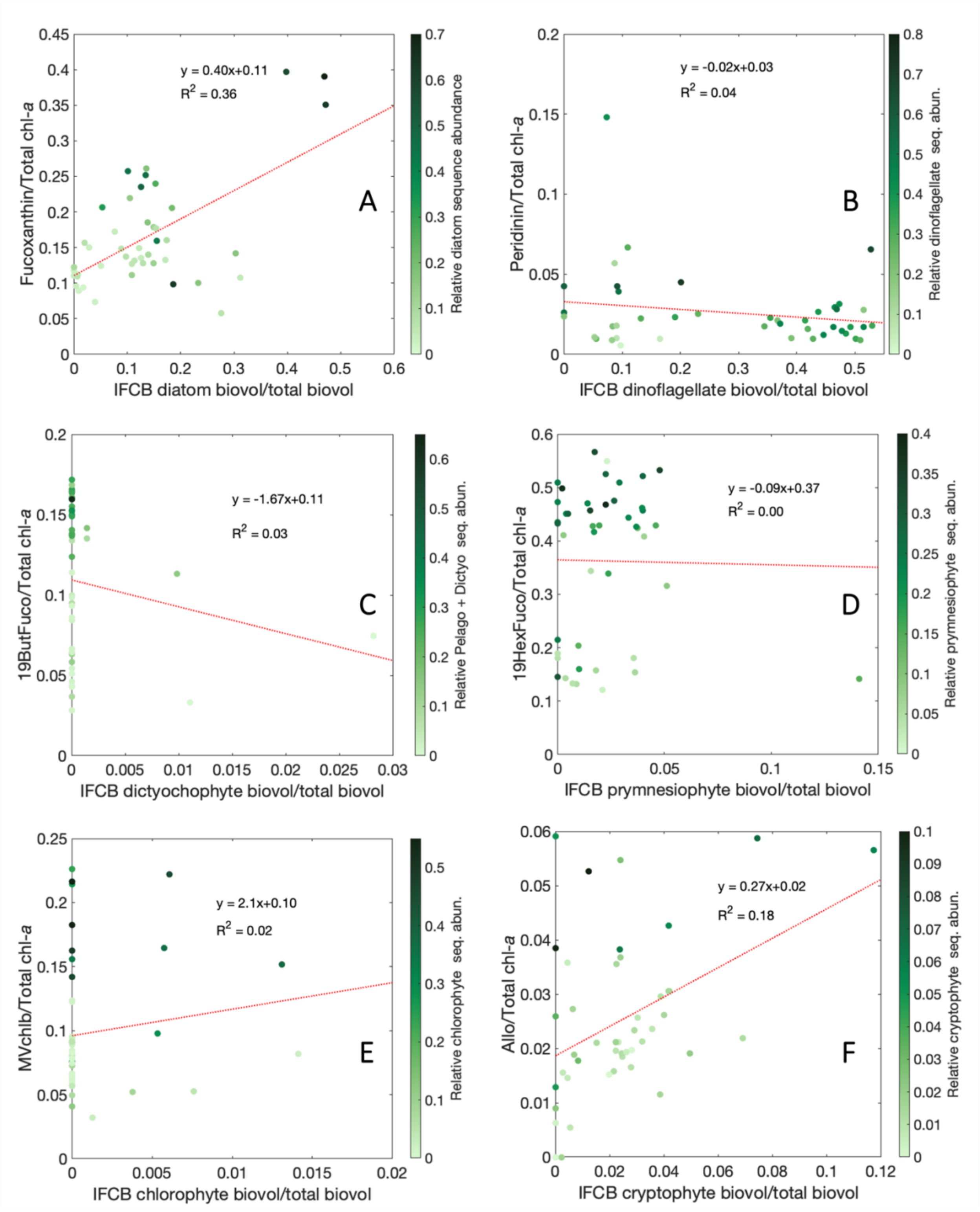

**Figure S3.** Relationships between relative pigment concentrations (normalized to Tchla) and relative biovolume fractions for (A) Fuco and diatoms, (B) Perid and dinoflagellates, (C) 19ButFuco and dictyochophytes, (D) 19HexFuco and prymnesiophytes, (E) MVchlb and chlorophytes, and (F) Allo and cryptophytes. All samples are colored by the relative fraction of 18S sequence abundances for the corresponding group.

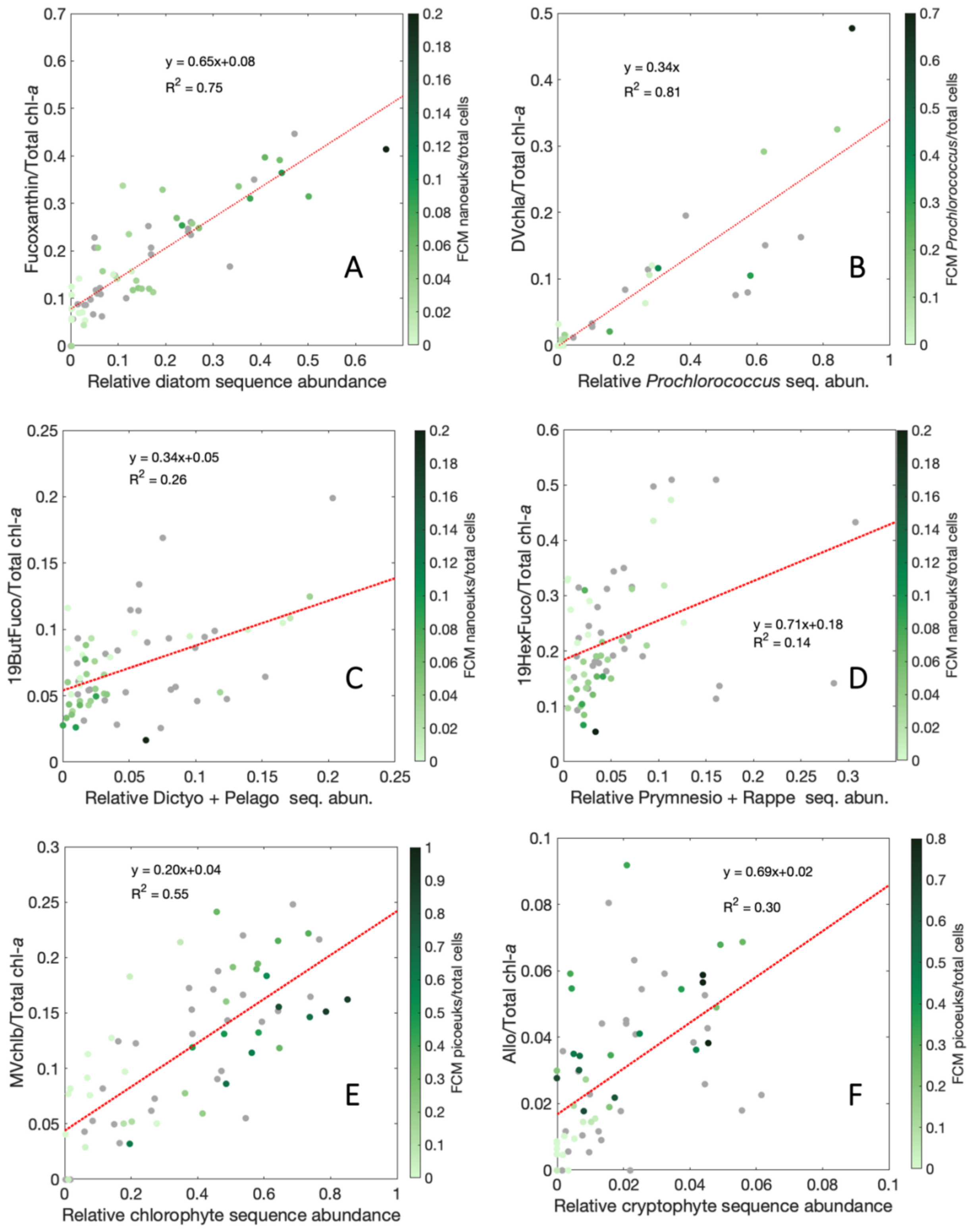


**Figure S4.** Relationships between relative pigment concentrations and relative sequence abundances for (A) Fuco and diatoms, (B) 19HexFuco and prymnesiophytes plus rappemonads, (C) Allo and cryptophyes, (D) 19ButFuco and dictyochophytes plus pelagophytes, (E) MVchlb and chlorophytes, and (F) DVchla and *Prochlorococcus*. All samples are colored by the relative FCM biovolume concentration that was determined to be most appropriate for that phytoplankton group. Gray dots represent samples for which there was not a FCM matchup.

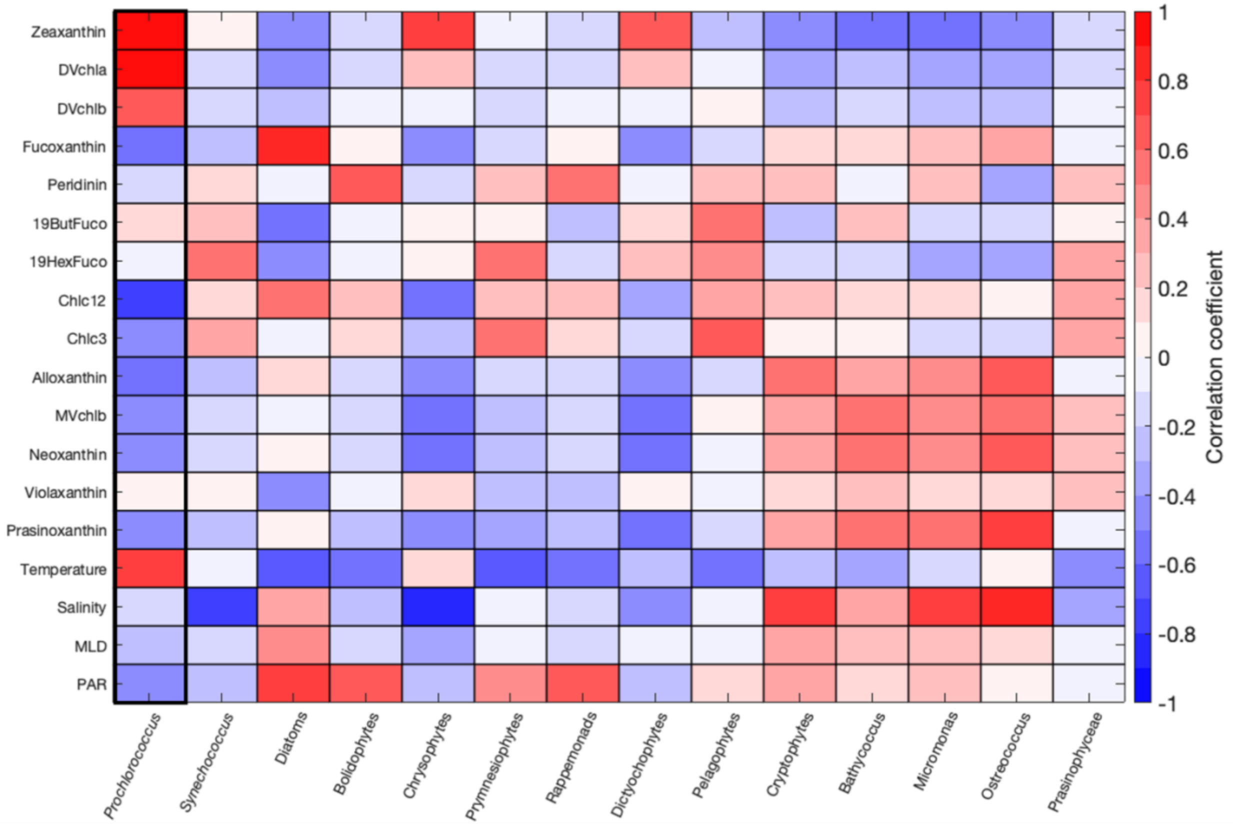


**Figure S5.** Pearson's correlation coefficient (R) between relative pigment concentrations and environmental variables (temperature, salinity, MLD, PAR) and relative sequence abundances from 16S.

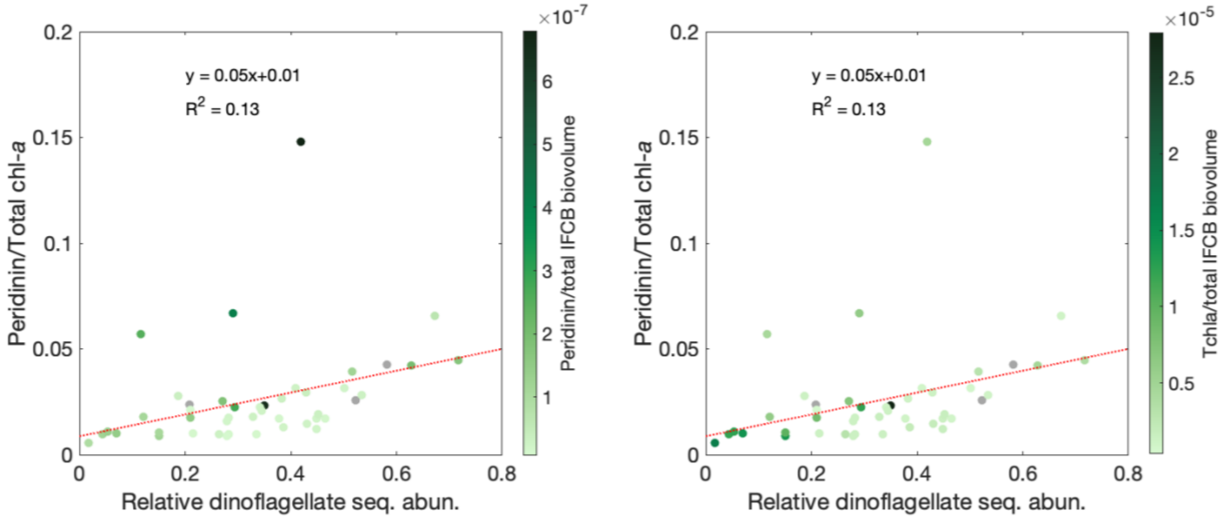


**Figure S6.** Perid/Tchla vs. relative dinoflagellate sequence abundance colored by (A) Perid concentration per IFCB biovolume concentration and (B) Tchla per IFCB biovolume concentration.
